# An Integrated Platform for High-Throughput Nanoscopy

**DOI:** 10.1101/606954

**Authors:** Andrew E S Barentine, Yu Lin, Edward M Courvan, Phylicia Kidd, Miao Liu, Leonhard Balduf, Timy Phan, Felix Rivera-Molina, Michael R Grace, Zach Marin, Mark Lessard, Juliana Rios Chen, Siyuan Wang, Karla M Neugebauer, Joerg Bewersdorf, David Baddeley

## Abstract

Diffraction-unlimited single-molecule techniques like STORM and (F)PALM enable three-dimensional (3D) fluorescence imaging at tens of nanometer resolution and are invaluable to investigate sub-cellular organization. The multitude of camera frames required to reconstruct a super-resolved image limits the typical throughput of these techniques to tens of cells per day, rendering these methods incompatible with large-scale cell biological or clinical application. STORM acquisition rates can be increased by over an order of magnitude, however the data volumes of about 40 TB a day and concomitant analysis burdens exceed the capacity of existing workflows. Here we present an integrated platform which transforms SMLM from a trick-pony technique into a work horse for cell biology. We leverage our developments in microscopy-specific data compression, distributed storage, and distributed analysis to automatically perform real-time localization analysis, which enable SMLM at throughputs of 10,000 cells a day. We implemented these advances in a fully-integrated environment that supports a highly-flexible architecture for distributed analysis, enabling quickly- and graphically-reconfigurable analyses to be automatically initiated from the microscope during acquisition, remotely executed, and even feedback and queue new acquisition tasks on the microscope. We demonstrate the utility of this framework by imaging hundreds of cells per well in multi-well sample formats. Our platform, the PYthon-Microscopy Environment (PYME), is easily configurable for hardware control on custom microscopes, and includes a plugin framework so users can readily extend all components of their imaging, visualization, and analysis pipeline. PYME is cross-platform, open source, and efficiently puts high-caliber visualization and analysis tools into the hands of both microscope developers and users.

---

Super-resolution Single-Molecule Localization Microscopy (SMLM) offers a roughly 10-fold improvement in resolution over conventional, diffraction-limited fluorescence microscopy, but it does so at the expense of acquisition time, data volume, and analysis overhead [1]. For SMLM techniques such as (d)STORM/(F)PALM, a single region of interest (ROI) a few ten microns in diameter usually requires a series of 10,000 to 100,000 camera raw data frames. These are typically acquired at 50 frames per second (FPS), meaning that a day of diligent manual imaging yields only a few tens of fields of view (Figure 1a). As a result, the majority of SMLM applications have addressed biological questions that can be answered with just a handful of super-resolved images, and have steered clear of questions requiring large sample sizes. A few rare and impressive quantitative investigations of phenomena that required sample sizes of hundreds of cells have been performed, but these required extensive data collection efforts spanning months to years [2]. Efforts to automate SMLM image acquisition [3–5] have reduced the amount of time the operator needs to spend in front of the microscope, relieving some of the tedium of acquiring large volumes of data, but have only resulted in modest improvements in overall throughput.

**Figure 1:**
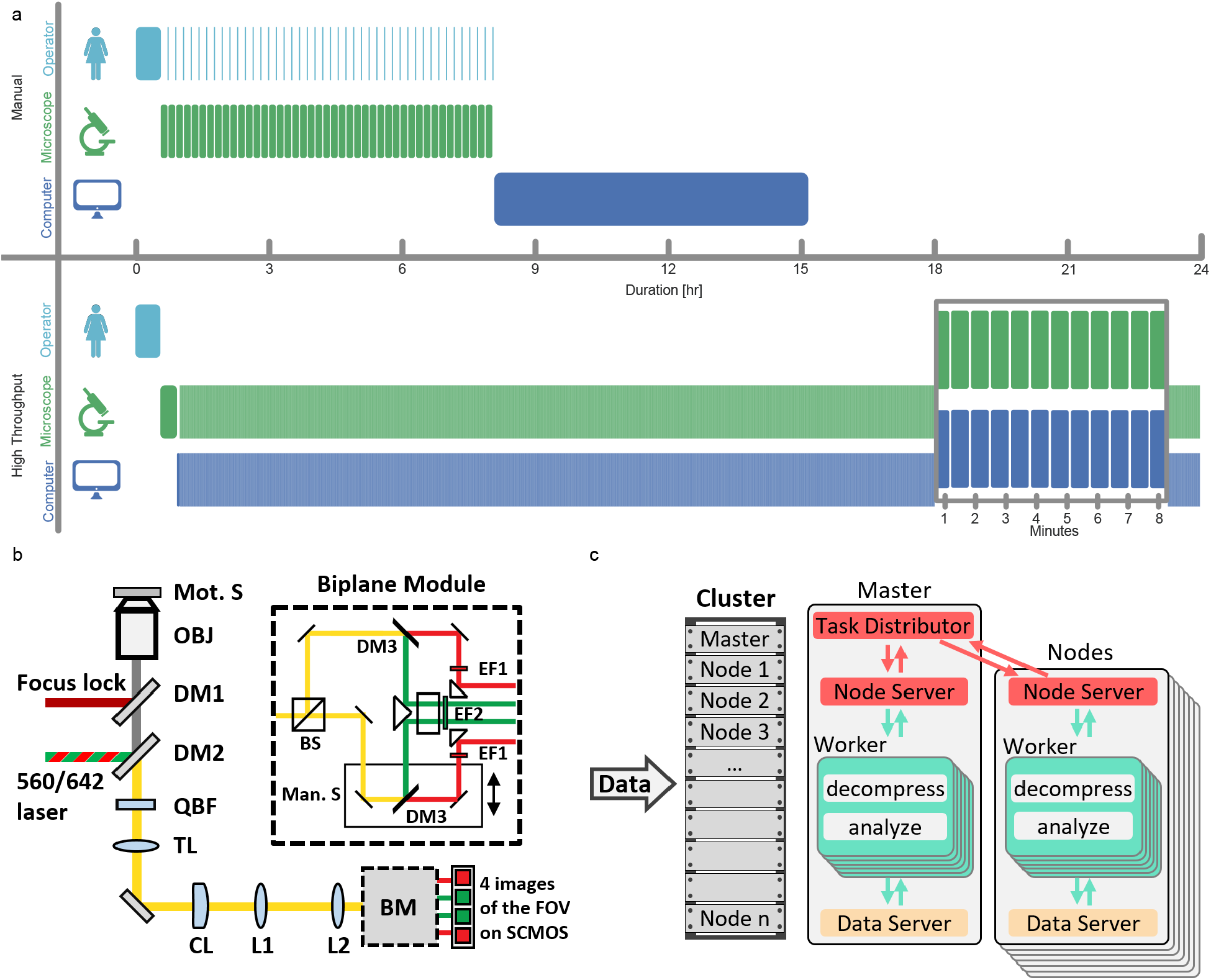
High-throughput SMLM. (a) Example timelines for SMLM acquisition of 36,000-frame ROIs performed at 50 FPS manually and automated at 800 FPS. (b) Schematic of automated multicolor 3D biplanar-astigmatism SMLM microscope. Mot. S.: motorized sample stage; OBJ: objective; DM1-3: dichroic mirrors; QBF: quad-band filter; TL: tube lens; CL: cylindrical lens; L1-2: relay lenses; BM: biplane module; BS: beamsplitter cube; Man. S.: manual translation stage; EF1-2: emission filters. (c) Diagram of scalable data pipeline for real-time localization and automated post-localization analysis.

By using sCMOS cameras [6] and high laser intensities it is possible to acquire SMLM data an order of magnitude faster [7, 8] than the typical 50 FPS, without a major loss in data quality. However, automating SMLM imaging at these high frame rates generates data at a rate of 800 Mb/s (70 TB/day) posing unique challenges for both data storage and analysis. The analysis burden is further compounded by the need to account for sCMOS-specific noise characteristics (which are more complex than those of EMCCDs [6]) in order to obtain high quality localization data. As a result, fast (> 400 FPS) SMLM imaging, has been largely restricted to 2D and has entailed a number of manual steps, both in image acquisition and analysis [6, 7, 9]. To deliver a truly high-throughput automated SMLM platform operating at sCMOS speeds represents a significant technological challenge requiring advances across microscope hardware, data handling, and analysis routines.

Here we present an integrated high-throughput SMLM platform operating at sCMOS speeds, transforming SMLM from an imaging technique specialized for small sample sizes into a high-throughput quantitative tool. We leverage our developments in data compression, distributed storage, and distributed analysis to automatically perform real-time localization analysis, and additionally present a flexible architecture for distributed and automatic post-localization analysis and feedback-based imaging workflows. Our multicolor 3D SMLM system is capable of imaging 10,000 mammalian cells a day, or entire studies configured on multiwell plates. The microscope control, data storage, and analysis pipeline integration is readily accessible to the community through the open-source PYthon-Microscopy Environment (PYME) [10], which additionally features advanced visualization [11] and plugin extensibility, making it a viable tool for complete and customized SMLM workflows.

## Results

### Imaging system

We built a microscope with hardware optimized for automated high-speed single-molecule imaging. It features an sCMOS camera capable of capturing a 2048×256 pixel region of interest at 800 Hz, high-power lasers, motorized lateral and axial stages, and a focus stabilization system. Custom spectral and focal splitting optics (Figure 1b, Supplementary Figure 1 and Supplementary Note 1.1) allow us to image two spectral channels, each at two different focal planes, simultaneously. By increasing the offset between focal planes from a typical biplane configuration [12, 13] to 750 nm and adding astigmatism [14] we can achieve high-quality 3D localization of single molecules over an extended axial range of about 1.2 μm. This halves the number of axial steps which need to be taken when performing 3D volumetric imaging, resulting in a corresponding improvement in speed. Using this setup, we can acquire multicolor 3D super-resolved images in about 10 seconds, as shown with several examples in Figure 2 (see also Supplementary Figure 2).

**Figure 2:**
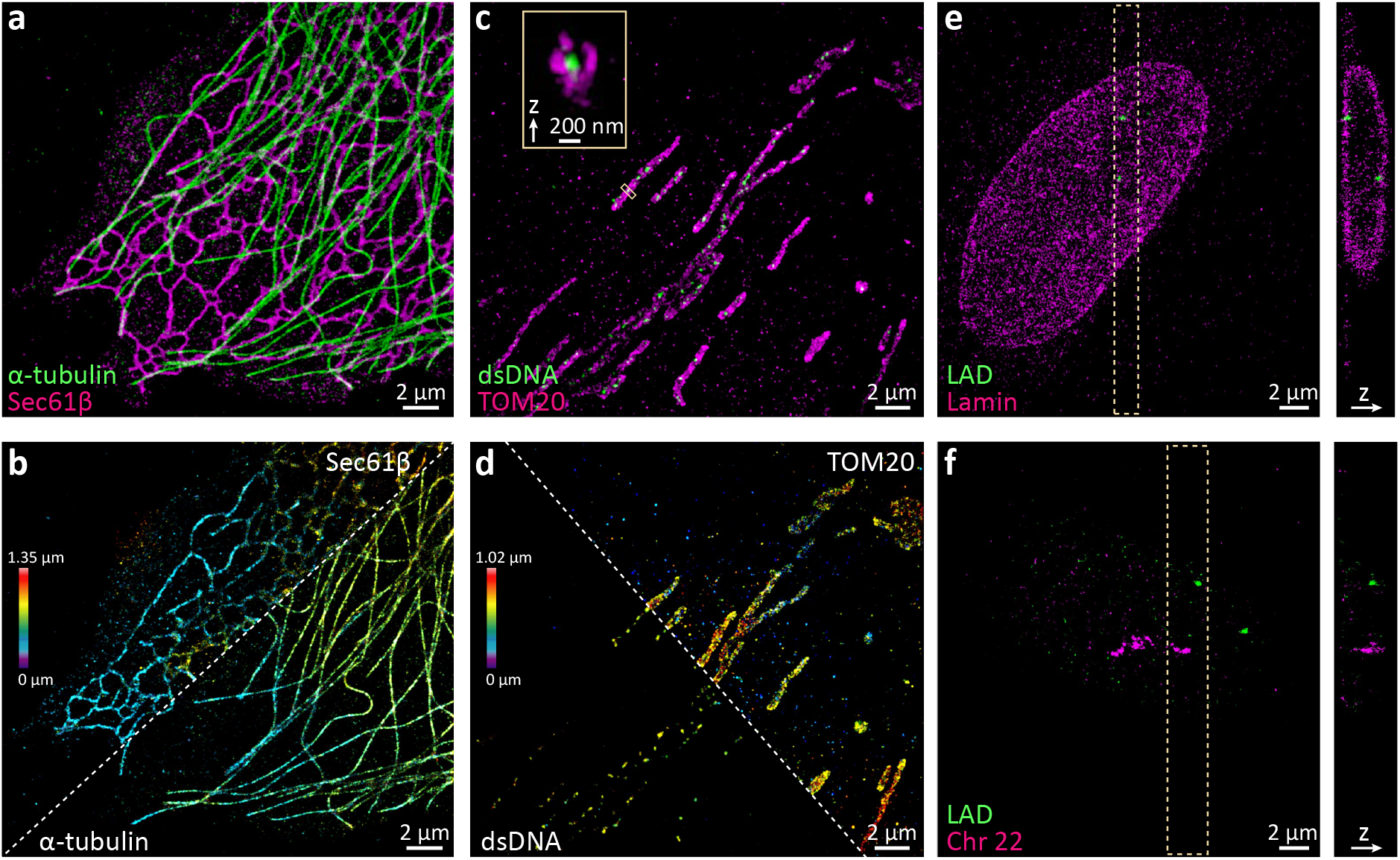
3D Multicolor Acquisition at 800 Hz Framerate (a, b) Rapid 2-color 3D SMLM of microtubules (alpha-tubulin immunolabeled with CF568) and endoplasmic reticulum (Sec61β-GFP immunolabeled with AF647) in a COS-7 cell. (c, d) Mitochondria (TOM20 immunolabeled with AF647) and nucleoids (doublestranded DNA immunolabeled with CF568ST) in a U-2 OS cell. (e) Lamin a/c (immunolabeled with CF568) and a lamin-associated chromatin domain (LAD, Chr13: 24405079-24709084, labeled with AF647 via FISH) in an IMR-90 cell. (f) All 27 topologically associating chromatin domains (TADs) along chromosome 22 (FISH-labeled with CF568) and a LAD (Chr5:115508197-115813276, FISH-labeled with AF647) in an IMR-90 cell. All data sets were acquired at a frame rate of 800 Hz for 8,000 frames (c-f) or 24,000 frames (g, h). Images are colored by label (a, c, e, f) or axial position (b, d).

### Scalable data handling - Compression, streaming, and distributed storage

Imaging at the full frame rate of an sCMOS camera generates an enormous amount of data (800 MB/s) and sustained imaging at this data rate is non-trivial. Previous high-speed SMLM efforts have saved data directly to a local solid-state drive (SSD) [6], but this limits acquisitions to a few hours before the SSD is full - even on the largest SSDs currently available. Once full, copying the data to slower storage (e.g. hard disk drives [HDDs]) can take significantly longer than the acquisition itself. To address the data movement and storage bottlenecks imposed by our high data rate, we developed a compression algorithm optimized for our microscope’s noise model with distributed storage across a small computer cluster (Figure 3a, Supplementary Note 3).

**Figure 3:**
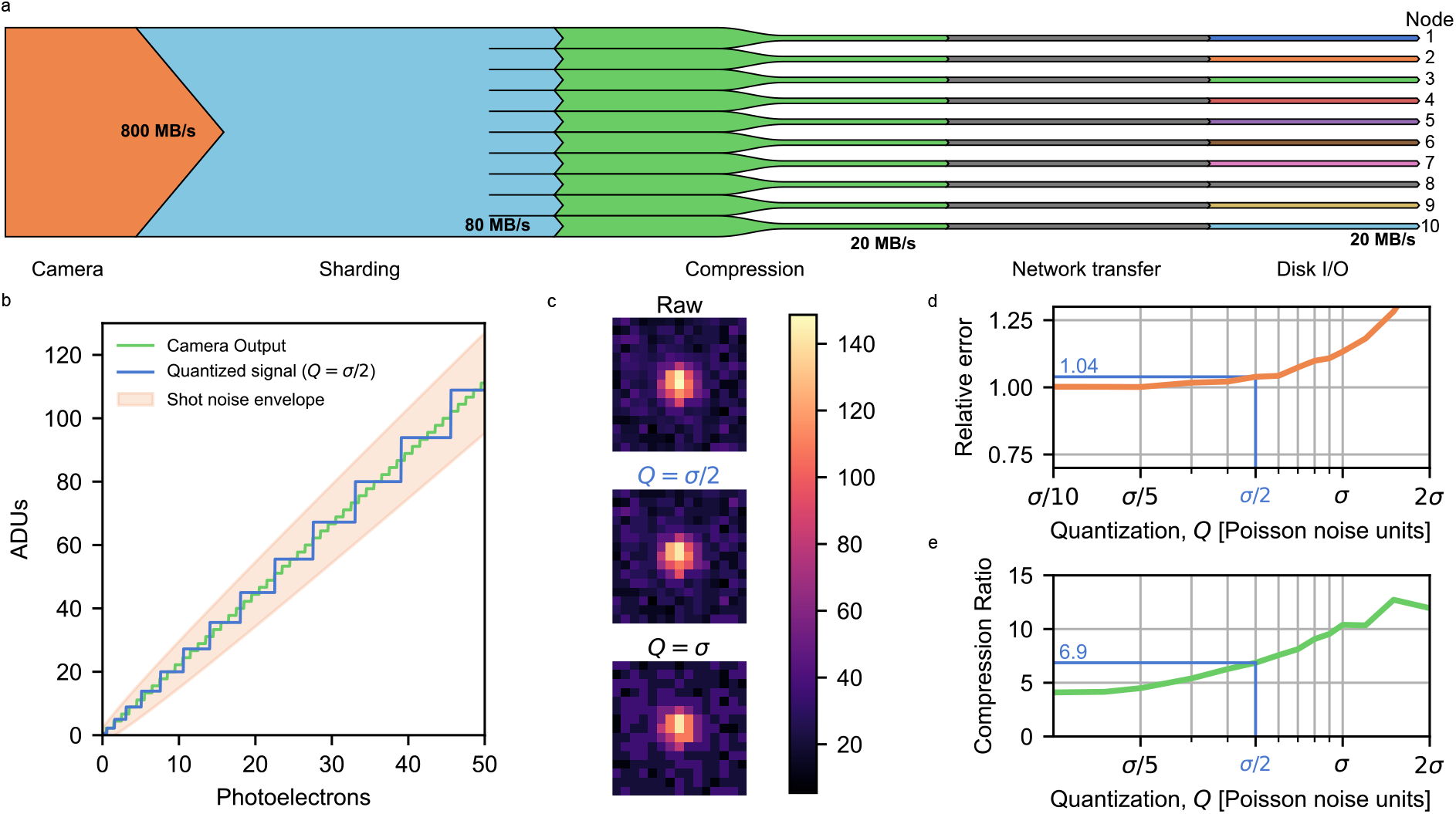
Data Volume Solutions. (a) Sankey diagram showing approximate data bandwidths as they are transferred from the camera to instrument computer RAM before being sharded, compressed (lossy), sent across a local network, and saved locally on hard disk drives on multiple computer nodes. (b) Our lossy compression algorithm re-scales the analog-digital units such that the corresponding number of photoelectrons represented by each unique value scales with a set fraction of the shot noise. (c) A simulated localization ROI shown at various quantization levels. (d) The relative localization error as a function of quantization for simulated localizations with an sCMOS noise model. (e) The compression ratio achieved at the same quantization levels as in (d).

Standard lossless compression algorithms such as zip offer a modest 2-3 fold reduction in file size when applied to SMLM raw data and are typically not fast enough to allow real-time compression at 800 MB/s. These algorithms use entropy coding (e.g. Huffman coding) which looks at the histogram of the data and uses short codes for frequently encountered values and longer codes for less frequently encountered values, combined with algorithms which encode repeated patterns. The poor compression ratios can be explained by two factors: SMLM raw data contain little repetitive structure for compression algorithms to exploit, in part due to Poisson noise, and the data as it comes from the camera is very conservatively quantized with more unique values than necessary to accurately represent the data; data from sCMOS cameras is typically quantized such that one photoelectron corresponds to ~ 2.5 analog-to-digital units (ADUs). This is reasonable for low signals (1-2 photoelectrons) where Poisson noise (which scales as 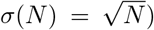 is ~ ±1 photoelectron (or ~ ±2.5 ADUs). However, a signal of 100 photoelectrons will have an error of ±10 photoelectrons (~ ±25 ADUs), giving rise to a band of 50 unique values which are not meaningfully distinct but will nonetheless make the signal harder to compress. The solution is simple: rather than using a constant quantization interval as provided by the camera, we re-quantize our data prior to compression such that the interval between quantal units is a constant fraction, *Q,* of the expected Poisson noise rather than a constant number of ADUs (depicted in Figure 3b).

We systematically varied *Q* for simulated SMLM raw data (Figure 3c, see also Supplementary Note 2) and analyzed the resulting relative localization error for obtained localizations (Figure 3d) and the achieved data compression ratio (Figure 3e). At *Q* = *σ*/2, we achieve 6.9-fold compression at a relative localization error of only 4%. This > 5 compression ratio holds across a large range of emitter densities (See Supplementary Figure 3).

While this re-quantization is technically lossy, it ensures that any losses are within the original data noise envelope (Figure 3b) and also visually preserves the integrity of blinking emitters, as seen in Figure 3c. Combined with the low relative localization error, we therefore deemed this to be an acceptable compromise and used *Q* = *σ*/2 in our further experiments.

Noting that the repeated pattern encoding portion of compression algorithms is not very helpful for our data, we skip this step entirely, greatly improving our speed (Supplementary Figure 4). The resulting compression is performed in real-time on the microscope computer and spooled over the network to a small computer cluster where each node is running PYME server processes (Figure 1c and Supplementary Figure 5). By sharding the data across multiple nodes on a per-frame basis, we further decrease the amount of per-disk bandwidth, and can use HDDs instead of SSDs for a low-cost, high-volume storage solution. This combination of noise-model aware compression and distributed storage thereby enables us to stream continuously at the full camera frame rate and also lays the foundation for analyzing the data in parallel.

### Real-time localization

Waiting for hours or days to do quality control or analysis on localization results largely defeats the purpose of imaging at high bandwidths. Although some simplified methods (such as centroiding) have been shown to yield high localization speed, these entail serious compromises in precision and accuracy [15]. Obtaining optimal localization precision with sCMOS cameras requires fitting using an sCMOS-specific noise model [6] and an algorithm such as Weighted Least Square [16] or Maximum Likelihood Estimation (MLE) [17].

We optimized our previous graphical processing unit (GPU) based MLE code [6] to achieve about 15-fold improvement in speed by using one thread per localization ROI pixel to evaluate the model function, with a single thread performing parameter updates on each iteration, rather than a single thread doing the entire fit (see Supplementary Figure 6). We also accelerated sliding-window background estimation and candidate molecule detection by moving them from the CPU to the GPU, highly parallelized using one-thread per x,y, (t) pixel (see Supplementary Note 5). Our now entirely GPU-accelerated localization pipeline performs about 10-fold faster than our CPU-only pipeline on a single computer (see Supplementary Table 1). However, even parallelized over multiple workers a single computer was too slow to keep up with our imaging.

Leveraging our distributed data storage, we developed a task-distribution architecture which enables a multiprocessing ‘cluster of one’ on a single computer as well as multi-computer clusters. Distributing tasks with a preference to assign jobs to computers where the data is saved allows us to minimize network overhead within the cluster. Critically, the performance of our architecture scales approximately linearly, allowing one to tune their localization speed simply by adding more computers. Our production cluster consists of ten computers made from affordable consumer-grade components in 2016; each equipped with a GPU to run our accelerated algorithm. This additional factor of ~10 improvement in performance allows us to localize in real-time (see Supplementary Table 1).

Localization tasks are automatically posted to the cluster on completion of recording or continually posted live during series acquisition for live visualization. A signal-to-noise based candidate molecule detection threshold enables localization to be performed automatically across a wide range of conditions without user attention.

### 3D Multicolor SMLM at 10,000 cells a day

The combination of our hardware and analysis advances enables us to image not only individual FOVs at high speed as shown in Figure 2, but 10,000 FOVs in a single day. To test this imaging mode, we plated U-2 OS cells on a coverslip and immuno-labeled their nuclear lamina (anti-lamin b1, AF647) and nucleoli (anti-NPM1, CF568). Automated imaging begins by first scanning the coverslip in widefield mode and stitching together the images to create a large mosaic image. This image is then segmented and processed to generate a list of suitable FOVs for SMLM imaging. The overview mosaic shown in Figure 4a was acquired in 52 minutes, after which automated super-resolution imaging of the 11,160 detected nuclei commenced. Each detected nucleus was imaged in 9.44 seconds, with the objective piezo actuator stepped over an axial range of 4.4 μm for an axial localization range of approximately 5 μm. The total imaging time for all 11,160 cells targeted was 1.2 days. 99.8% of target FOVs were successfully acquired (see Methods), resulting in a total of 3,589,123,170 fitted emitters from 75,431,069 raw frames (504,879,862 high-precision localizations after combining molecules present in more than one of the biplane views and filtering). The first and last nuclei imaged are shown in Figure 4b. While they look relatively similar to each other, we also observed oddly shaped nuclei and nuclei which one could reasonably think were representative if only imaging a handful of cells. To demonstrate the latter, we performed a principal component analysis (PCA) on a collection of features extracted from the SMLM localization data (see Supplementary Table 2 and Supplementary Figure 7). We show the closest cell to the PCA-space mean position (Figure 4c), and cells located two median absolute deviations away along both of the principle axes (Figure 4d-g). Figure 4h shows the ensemble median-normalized NPM1-NPM1 pairwise distances of these selected cells, which vary substantially outside the interquartile range depicted by the gray-shaded area.

**Figure 4:**
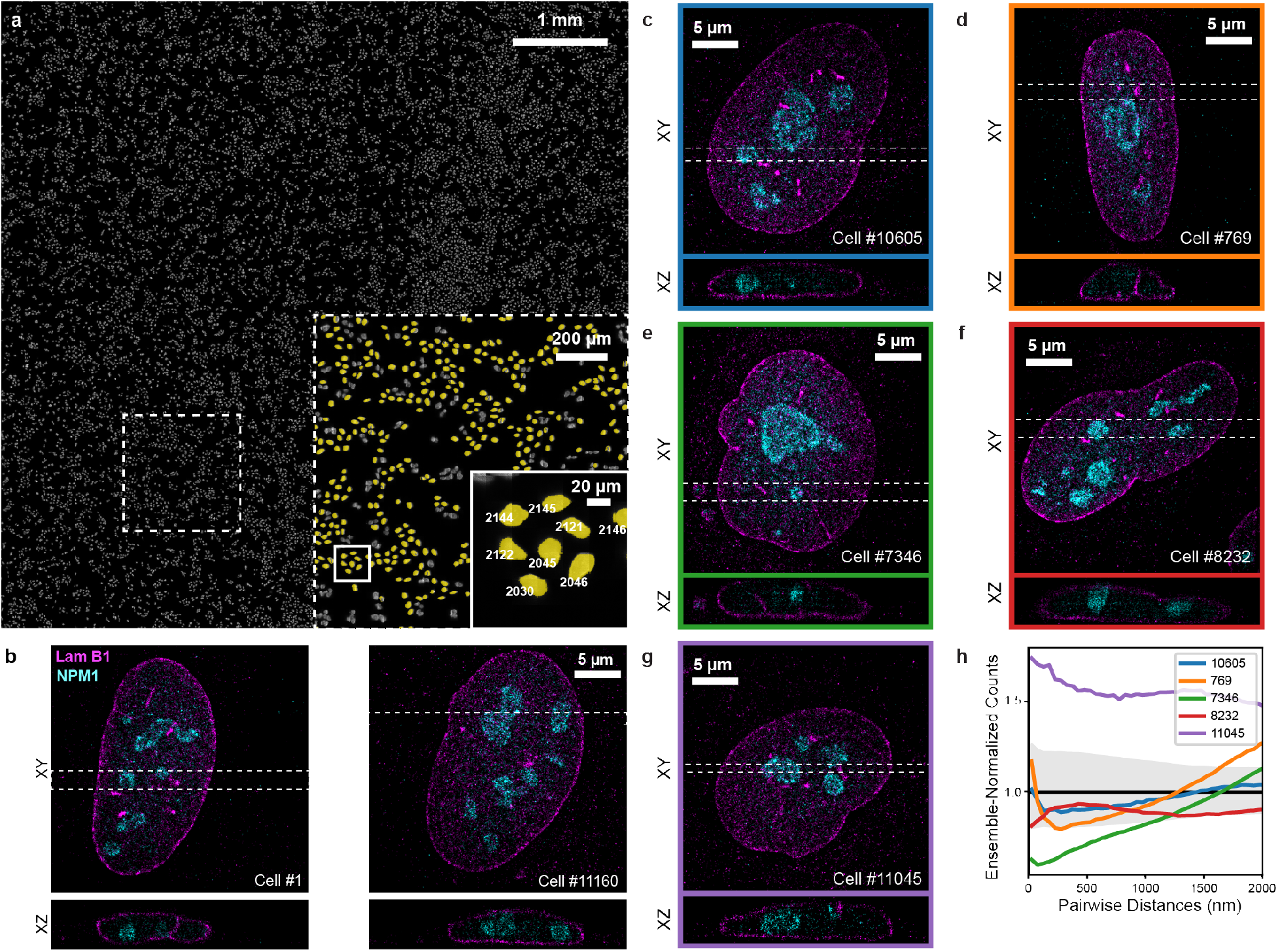
3D Multicolor SMLM imaging of 10,000 cells a day. (a) Overview mosaic image of U-2 OS cells on a coverslip, from which 11,160 fields of view were automatically detected for imaging. A magnified view of the dashed box in the overview image is shown in the large inset, with a further magnification shown inside the solid box. Nuclei that were queued and imaged are highlighted in yellow and their queue number displayed in the smaller inset. Each detected nucleus was automatically imaged, averaging 9.44 seconds per FOV, or 10,000 cells per 26.2 hours. The first and last nucleus imaged are shown in (b). Principle component analysis on the SMLM datasets was used to select representative cells, choosing the cell closest to the mean (c, blue) and ±2 median absolute deviations from it (d-g). (h) Sum-normalized nucleophosmin-nucleophosmin pairwise distance histograms for these selected cells c-g are shown after a secondary normalization to the ensemble-median (black, interquartile range shown in gray).

### Flexible analysis and integrated workflows

Analyzing thousands of SMLM localization datasets has to date been non-trivial, and often relied on patching together multiple existing packages. The analysis backbone of PYME, ‘recipes’, can be graphically reconfigured, handle hybrid pointcloud/image-data workflows, and can be quickly built and tested in PYME’s interactive viewers [11]. Recipes can be efficiently batched on a single computer, or run on the cluster using the same analysis distribution we employ for localization tasks. They can additionally be chained together, allowing automatic localization and post-localization analyses workflows.

Microscope control solutions for custom systems are often bespoke, with their control flows hard-coded, making maintenance and code sharing difficult. Many automated control flows consist of image analysis steps, particularly for more intelligent automation, which can quickly compound the amount of special-casing in instrument control code if these become at all sample-specific. PYMEAcquire, used here, allows intricate and easily re-configurable workflows by using a priority queue for acquisition tasks. Acquisition protocols can be quickly written as lists of tasks to be executed on specified frames. Analysis chains can then be built graphically and linked to specific protocols, triggering (remote) execution on the PYME cluster at the beginning or end of a series acquisition as set by the user (see Supplementary Note 4). Finally, server endpoints in PYMEAcquire allow remote queuing of acquisition tasks, and can be leveraged in recipe modules to add acquisition tasks to the priority queue based on analysis of previously acquired series. This establishes a flexible and powerful architecture, fully integrating the instrument with distributed analysis.

### Localization microscopy meets high-content screening

Conducting entire multi-condition studies by SMLM has until now been impractical. Even in automated workflows, low frame rates coupled with acidifying glucose-oxidase/-catalase STORM buffers have limited the total number of ROIs imaged per plate to less than 100 [4]. We leveraged our platform to study the distribution and size of Cajal bodies (CBs), the site of small non-coding RNA transcription and processing, in cells undergoing osmotic shock [18, 19]. Cellular stresses disrupt gene transcription, and as many nuclear bodies are hypothesized to be nucleated by transcription of specific genes, we expect the global disruption of transcription and RNA synthesis induced by osmotic shock to produce profound effects on Cajal body integrity [20].

We imaged HeLa cells immuno-labeled for coilin, a Cajal body mark, with Cy3b, and lamin b1 with CF660C in an 8-well slide format. A tile-overview in the lamin channel was queued for each well, and we chained the overview protocol to a nucleus detection recipe to queue 45,000-frame 800 Hz SMLM series at a maximum of 75 ROIs per well in a path-optimized manner and at a higher priority than the tile overview tasks such that large movements across the slide were minimized. We observed relatively consistent presence of Cajal bodies per cell in the control condition (median, mean, std. dev. = 1, 1.4, 1.1, N=71. See Supplementary Figure 8 for a histogram of number of CBs per ROI). An ROI automatically selected as representative of each well by PCA is shown in Figure 5a (see Methods). The CBs were indeed profoundly affected, and were depleted in wells which had been subjected to substantial osmotic shock, with no CBs detectable at concentrations higher than 75 mM. Additionally, the CBs decreased in size with increasing osmolarity (see Figure 5c).

**Figure 5:**
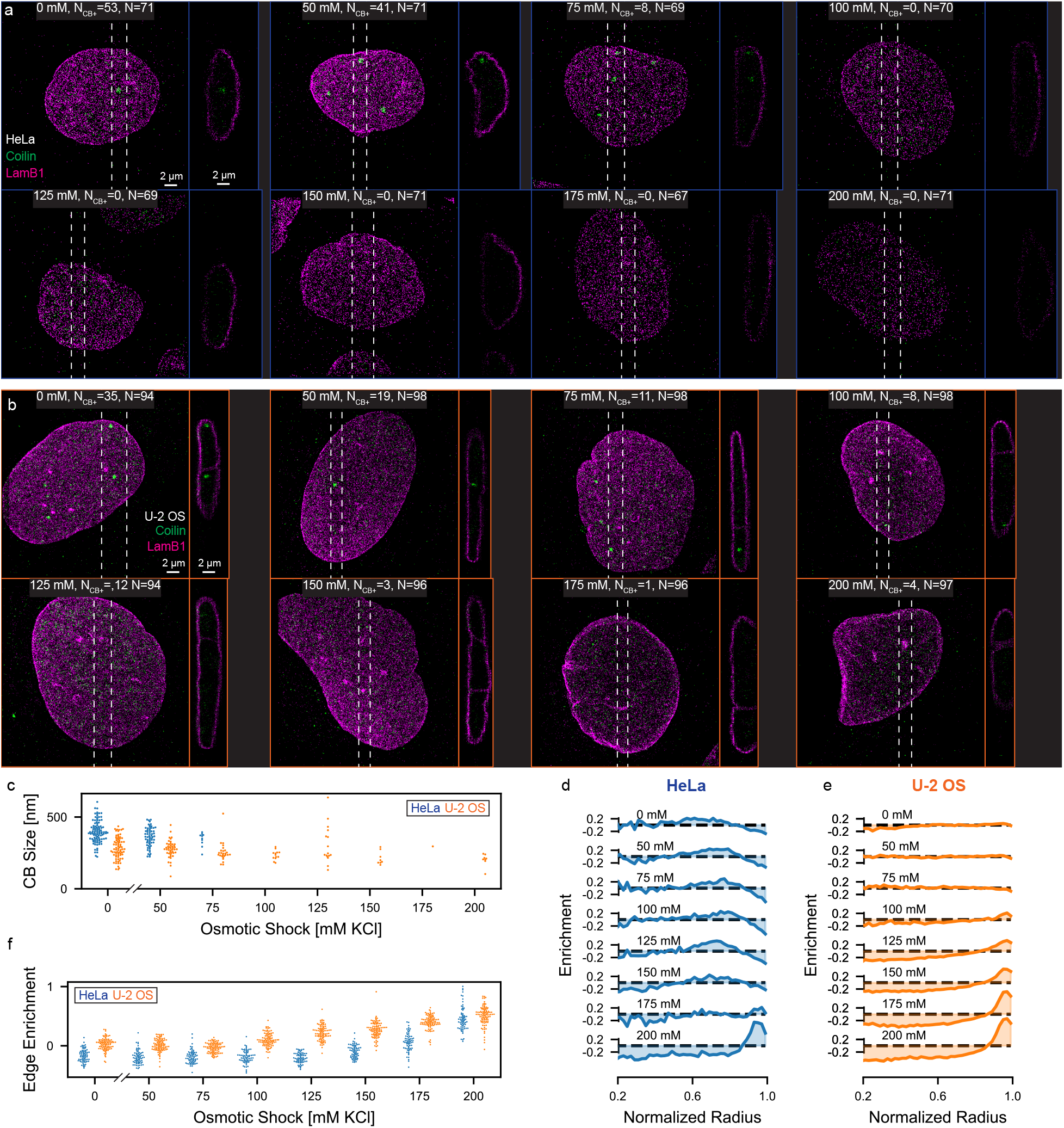
(a, b) Example PCA-selected ROI from an osmotic shock experiment on HeLa cells (a) and U-2 OS cells (b) in 8-well slides. The concentation of KCl, number of ROIs containing a segmented Cajal body (N_CB+_), and number of ROIs successfully imaged and analyzed (N) are annotated on the example ROI for each condition. (c) Size of CBs at each concentration. (d, e) Coilin enrichment relative to a uniform random distribution within a fitted nucleus model for HeLa cells (d) and U-2 OS cells (e). (f) Coilin edge enrichment for each ROI, calculated as the average coilin enrichment relative to a uniform random distribution at normalized radii larger than 0.85.

To see if this was an idiosyncrasy of the cell type or a more universal phenomenon, we repeated the experiment in U-2 OS cells, imaging a maximum of 100 ROIs in each well for 33,000 frames each (Figure 5b). Notably, the CBs in the U-2 OS control cells are smaller, and their number more varied (median, mean, std. dev. = 0, 0.8, 1.4, N=94; see also histograms in Supplementary Figure 8), making them generally more difficult to study at low throughputs. Further, while the osmotic shock still affects both CB presence and size, these effects are more subtle than in the HeLa cells (Figure 5c).

While the dissipation of Cajal bodies under osmotic shock could be rigorously monitored with our system, the function of most proteins in membraneless organelles is not limited to these structures, and we noticed an opportunity to leverage our statistical power to measure the redistribution of coilin. For each ROI, we fit the lamin localizations to a spherical harmonic shell in order to define a nuclear coordinate system for the Coilin localizations. We see Coilin enrichment (relative to uniform distribution simulated in the shell) at the nuclear periphery in stressed HeLa and U-2 OS cells (Figure 5d-f).

## Discussion

We have shown that it is possible to carry out large SMLM studies without sacrificing sample population or experimental conditions. This is a substantial departure from the current state-of-the-art. The software advances this required are not only enabling of higher-throughput studies, but should additionally reduce frustrations common with smaller experiments or even manual exploration. The modular and plugin-friendly nature of PYME and its interactive data viewers make it feasible for users to easily explore and extract user-defined features from these large data sets. Similarly, PYME is well suited for instrument developers to easily extend their acquisition or analysis capabilities in a complete environment.

Automated SMLM at high bandwidths enables even casual users to acquire more data than singlemolecule imaging experts, which, while quite powerful, additionally warrants consideration of tools to assist them. For example, localization quality control such as Wasserstein-induced Flux [21] and ROI error-mapping methods such as SQUIRREL and HAWKMAN [22, 23] could be helpful in determining if imaging and localization analysis were performed adequately. PYME recipes can produce HTML report outputs, and plugins to calculate such measures of quality control could facilitate automatic checks for each series or acquisition run.

In addition to monitoring image quality, our approach could be improved by further optimizing automated acquisition, for example: regulating emitter density by servo-controlled laser intensities, varying the number of frames acquired at a given ROI depending on its actual sampling requirements, or correcting sample-induced aberrations at each ROI. While impressive progress has been made in the area of intelligent SMLM automation [24], there are many unmet challenges to apply these advances to 3D and/or multicolor imaging, and especially for high-speed imaging at camera frame rates of 800 Hz.

We expect hardware advances such as higher-bandwidth cameras with reduced amplification noise and larger fields of view to continue to improve SMLM image quality and capabilities. We additionally note that per-pixel quantization using noise envelope scaling as demonstrated in this work could be performed on board cameras, and could aid in further bandwidth optimizations.

With the advances contained in this work, we expect SMLM to become a much more routinely and broadly applicable tool. We further anticipate that our developments will help bring large-scale tissue applications and even clinical use within reach.

## Methods

### High-Throughput SMLM Microscope

The optical setup, shown in Figure 2b is detailed in Supplementary Note 1.1 and Figure 1, and discussed here only briefly. The microscope hardware differs from conventional SMLM microscopes in that an sCMOS camera and two high-power (2W) excitation lasers (560 nm and 642 nm) are used. A cylindrical lens to provide astigmatism is combined with a biplane module to extend the axial localization range. The laser lines are coupled into a multimode fiber, which is vibrated to average out speckle and achieve a uniform intensity profile at the fiber exit which is imaged into the sample. A fast objective piezo actuator enables axial scanning, and a motorized sample stage accommodates lateral sample movement on the order of 100 mm. The microscope additionally has a custom-built focus lock, where a near-infrared laser is reflected off the coverslip at an angle such that the reflected beam position can be monitored by an additional camera. This position indicates the distance between the objective and the coverslip, and can be servo-controlled.

The instrument was originally controlled with a custom LabView program (Phase 1), which was used for imaging in Figures 2 and 4, and would save DCIMG formatted image files to a RAM disk where a PYME script would open them and spool them to the computer cluster, additionally launching localization analysis. We then converted instrument control to PYMEAcquire for further integration of the system (Phase 2), after which we no longer saved frames to disk on the instrument computer as we could compress and spool them directly.

### SMLM Imaging

Alexa Fluor 647 (AF647) and CF660C are suitable for high-speed SMLM [7, 8], CF660C being particularly robust. We additionally found CF568 to be capable of fast switching, enabling high-speed multicolor SMLM in standard STORM buffers using the glucose-oxidase/-catalase oxygen scavenging system (‘glox’ buffer). Not all imaging formats are easily sealed, and we find sulfite-based oxygen scavenging in aqueous buffer [25] performs well at high speeds, and is compatible with Cy3B and CF660C, again enabling sustained high-speed multicolor imaging.

The glox buffer used in this work consisted of 0.49 kU/mL glucose oxidase, 0.98 kU/mL glucose catalase, 0.14M 2-mercaptoethanol, 2.5mM Tris at pH 8, 2.5mM NaCl, and 0.28M Glucose in milli-q water. The aqueous sulfite buffer used in this work consisted of 20 mM Na_2_SO_3_ and 0.14 M 2-mercaptoethanol in 1X PBS.

### Sealed glox buffer imaging

All samples in Figures 2 and 4 were imaged in the glox STORM buffer. These samples were mounted and imaged in Bioptechs FCS2 flow chambers with the microaqueduct slide flipped upside down such that the flow input/output tubes were sealed. Before mounting the sample, the edges were scraped with a scalpel to remove cells which would interfere with ideal sealing. We used 0.5 mm thick circular gaskets, overfilled the chamber with 500 μL of STORM buffer and rinsed the coverslip in 200 μL of STORM buffer before mounting. Once the coverslip was in place, it was flattened (and excess buffer was removed) by pressing a wipe against it using a flat plastic dish.

All SMLM imaging shown in Figures 2 to 4 was performed at a camera frame rate of 800 Hz. The 560 nm and 642 nm laser intensities delivered to the samples were approximately 71 kW/cm^2^ and 51kW/cm^2^ for the lamin B1 and NPM1 sample, 41 kW/cm and 51 kW/cm for the lamin A/C and LAD pool 2 sample, 61 kW/cm^2^ and 51 kW/cm^2^ for the *α*-tubulin and Sec61*β* sample, 38 kW/cm^2^ and 51 kW/cm^2^ for the dsDNA and TOM20 sample, and 41 kW/cm^2^ and 61 kW/cm^2^ for the Chromosome 22 TADs and LAD pool 1, respectively. 405 nm activation light was not used during SMLM data acquisition of these samples.

The 3D images of the *α*-tubulin and Sec61*β* sample, and the TOM20 and dsDNA sample, were acquired without ‘z-stepping’ during recording. The remaining samples in Figures 2 and 4 were imaged with 2 or 6 cycles of 7 interwoven ‘up then down’ z-steps to obtain even axial localization distribution over the thicker volumes.

### Sulfite buffer imaging

The HeLa and U-2 OS samples in Figure 5 were imaged in an aqueous sulfite STORM buffer, which enables flexible sample formats such as small petri dishes, 8 well slides, and multi-well plates. For these samples, N_2_ was additionally perfused into a stagetop incubator chamber (OKO K-Frame) at a rate of 0.2L/min. Stage leveling was checked before imaging, and the offsets measured by the leveling routine were stored to bootstrap automatically finding the coverslip should the focus lock ever be lost.

All SMLM imaging with the sulfite buffer (the coilin and lamin samples in Figure 5) was performed at 783 Hz. The 560 nm and 642 nm intensities delivered to the samples were approximately 49 kW/cm and 53 kW/cm^2^. Imaging began on each ROI only with 642 nm illumination, turning on the 560 nm laser after a full z-stack cycle (18,000 frames for the 45,000 frame HeLa series, and 11,000 frames for the 33,000 U-2 OS series. Additionally <0.08 kW/cm^2^ of 405 nm illumination was applied nearer the end of each series. The U-2 OS series of 33,000 frames were acquired in about 44 seconds each.

### Compression

Prior to spooling the data from the instrument computer RAM to the computer cluster, we minimize the data volume via compression. This compression includes 3 steps: offset subtraction, re-quantization such that the quantization intervals scale with the square root of the number of photons, and then Huffman coding [26]. The re-quantization step serves to decrease the number of discrete levels within the data, dramatically increasing the efficiency of Huffman coding. This step at each j pixel can be expressed as

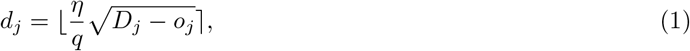

where *D* is the intensity counts in a single frame, o is the camera offset map, *d* is the resulting quantized intensity bin, *η* is the number of photoelectrons per analog-to-digital unit, and *q* is the desired scale factor in poisson noise units (i.e. 0.5 for *σ*/2 quantization). ⌊x⌉ denotes a rounding operation on *x* to the nearest integer.

Our approach achieves a 5-8 fold reduction in data volume. While the quantization step is lossy, we choose the quantization intervals such that the additional quantization noise introduced is significantly less than the Poisson noise we would expect from photon detection and therefore has a negligible influence on subsequent processing and localization. Our approach is similar to one developed separately and reported in [27]. Our optimized C code leverages the AVX SIMD instruction set when available and allows the compression to run at ~1 GB/s without the need for specialized hardware (e.g. GPUs) on the acquisition machine (see Supplementary Figure 4).

### Lamin and Nucleophosmin PCA

Of the 11,160 series imaged, 11,136 were spooled and localized during the same time, with analysis being run post-facto on an additional 9 series for a total of 11,145 SMLM images. Spooling failed for 15 images. Additionally, pointclouds with less than 5,000 localizations were ignored. For each of the remaining 11,117 lamin and nucleophosmin SMLM point clouds, 13 metrics were calculated and used to create a feature vector. These metrics are calculated directly from the localizations and are described in the Supplementary Table 2. The feature vector was normalized by the interquartile range along each metric, and this 11,117 × 13 array was then reduced to size 11,117 × 2 using principal component analysis (PCA) limited to two components. The features for each of these 11,117 point cloud were projected onto the resulting principle component 0 and 1 (PC0, PC1) 2D basis and plotted in Supplementary Figure 7. The median absolute deviation (MAD) was calculated along PC0 and PC1, as was the mean of the ensemble. A KDTree of PC0/PC1 euclidean distance was generated and the cells nearest to the mean, and the mean ±2 MAD along each axis were selected as shown in Supplementary Figure 7, and displayed in Figure 4c.

### Cajal body measurements and Coilin distribution analysis

Cajal bodies were segmented from coilin localizations using a Bayesian Information Criterion (BIC) optimized Gaussian Mixture, and lamin localizations were fitted to an expansion of spherical harmonics to create a radial reference frame for each coilin localization.

First, a generous 2D binary mask of the nucleus was used to isolate signal localization for all further analysis. This mask was generated from the lamin localizations by rendering a 2D *σ* = 30 nm Gaussian image with 40 nm pixels, and applying a 15 × 15 pixel maximum filter, followed by an Otsu threshold. Regions other than the largest were discarded, and the nucleus mask was dilated 1 pixel 10 times.

#### Radial Coilin distribution analysis

The lamin localizations were least squares fitted to an expansion of 16 spherical harmonic functions (the *l* ∈ {0, 1, 2, 3} modes) centered at the lamin localization center of mass, producing an analytic shell representation of the nuclear envelope. Coilin localizations were converted from Cartesian to spherical coordinates, again relative to the lamin center of mass, and the shell radius at their zenith and azimuth coordinate was used to normalize their radial coordinate. This shell-normalized Coilin distribution was histogrammed into 50 bins and sum-normalized. In order to compare the Coilin distribution with a uniform distribution, this process was repeated for a uniform-density point-cloud simulation bounded by the same shell (5 iterations targeting a density of 1 point per isotropic 200 nm voxel). Finally, a per-nucleus Coilin-enrichment histogram was created by dividing the shell-normalized Coilin radius histogram by that of the uniform-density simulation and subtracting 1. The per-well average of this Coilin-enrichment histogram is shown in Figure 5d, e. The per-nucleus edge enrichment (Figure 5f) was calculated by taking the average value of the Coilin-enrichment in radial bins above 0.85.

#### Cajal body measurements

A Gaussian Mixture Model (GMM) was fitted to the Coilin localizations. For each ROI, the number of components was stepped by 1 from 1 to 50, and the BIC was calculated for each. Metrics were calculated for each candidate CB as shown in Supplementary Table 3. These candidate CBs were then filtered, rejecting candidates with less than 20 localizations, greater than 711 nm MAD (corresponding to to a Gaussian FWHM of ~2.5 μm) and an approximate density of at least 5 × 10^-6^ nm^-3^ (with volume calculated as 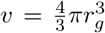, where *r_g_* is the gyration radius).

The ROIs shown in Figure 5a,b were selected using PCA of these per-CB features by well. For each well, if there were CBs in less than 5% of the ROIs the feature vector (see Supplementary Table 3) was constructed from the raw BIC-optimized GMM results (candidate CBs) rather than the filtered (CBs) as was otherwise used. After that decision, the feature vector was scaled by centering on the feature medians and normalizing by their interquartile ranges before performing a 3-component PCA. The feature vector was then projected onto these principle components, and the median value along each component was used to query a KDTree of PC0/PC1/PC2 euclidean distances. Each ROI shown in figure 5 contains either the closest or second closest CB candidate to that point.

The GMM implementation used is in Scikit-Learn, and each component was fitted with its own general covariance matrix [28].

## Code Availability

The code for distributed data storage and analysis is released under the GNU General Public License v3 as part of the python-microscopy project [10] Code for GPU acceleration of single-molecule fitting is available under an academic use license from github.com/barentine/pyme-warp-drive. The LabVIEW acquisition software used in phase 1 can be obtained from the authors, however is not actively maintained. Please contact the authors for alternative licensing arrangements.

## Data Availability

The 3D localizations, calibration files, and raw blinking movies for all series in Figure 2, and Cell #1, #2504, #5735, #8041, #9577, and #11,160 from the lamin-NPM1 dataset in Figure 4 (and 3D localizations for the remaining cells) are publicly available through the 4D Nucleome data portal. Any additional data from this work can be obtained through the authors upon request.

## Acknowledgements

We are grateful to Allison Wrogg for contributing the timeline in Figure 1. We thank Lena Schroeder and Yongdeng Zhang for helpful discussions and technical assistance. This work was primarily supported by a 4D Nucleome grant from the National Institutes of Health (U01 DA047734 to J.B. and D.B.). J.B. acknowledges support from NIH grant P30 DK045735 (to Robert Sherwin). A.E.S.B. acknowledges support by a NIH training grant (T32 GM008283) and training at the Computational Image Analysis in Cellular and Developmental Biology Course of the Marine Biology Laboratory (which was supported by NIH R25 GM103792).

## Author Contributions

Y.L. and J.B. designed the optical hardware of the microscope which Y.L. built. A.E.S.B., Y.L., D.B., T.P., and J.R.C. developed acquisition control software. M.R.G., A.E.S.B. and D.B. designed and implemented the computer cluster. D.B. designed the distributed storage architecture and compression algorithm. D.B. and L.B. designed and implemented the cluster task distribution. A.E.S.B. and D.B. developed the GPU acceleration code. S.W. and M.L. designed the FISH probes. E.C., P.K., M.L., F.R.M., M.D.L., S.W., and K.M.N. optimized sample preparation protocols and prepared samples. A.E.S.B., Y.L., and E.C. performed imaging experiments. A.E.S.B., Y.L. and D.B. performed post-localization analysis. All authors contributed to writing the manuscript.

