## Supplementary Information for "An Integrated Platform for High-Throughput Nanoscopy"

##### External Files

Supplementary Video 1: Video showing a reconstruction each of the 11,145 nuclei contributing to Figure 4.

Supplementary Extended Table 1: LAD probe design including sequences of PCR primers, reverse transcription primers, and activator oligos. The worksheet lists the sequences of the forward and reverse PCR primers and the reverse transcription primers used in the synthesis of each LAD probe library. The /5Alex647N/ in the sequences indicate 5' Alexa Fluor 647 modifications of the DNA oligos.

Supplementary Extended Table 2: Template sequences for LAD probe library syntheses. All oligonulceotide sequences in each template LAD probe library are compiled as a list. Different LAD libraries are placed in different worksheets labeled with the names of the LADs that the libraries target.

Supplementary Extended Table 3: TAD probe design information including sequences of PCR primers and reverse transcription primers. The worksheet lists the sequences of the forward and reverse PCR primers and the reverse transcription primers used in the synthesis of the Chr22 probe library. The /5Biosg/ in the sequence indicates 5' biotin modifications of the DNA oligos.

Supplementary Extended Table 4: Template sequences for TAD probe library syntheses. All oligonulceotide sequences in each template LAD probe library are compiled as a list. Different LAD libraries are placed in different worksheets labeled with the names of the LADs that the libraries target.

### Supplementary Figures and Tables

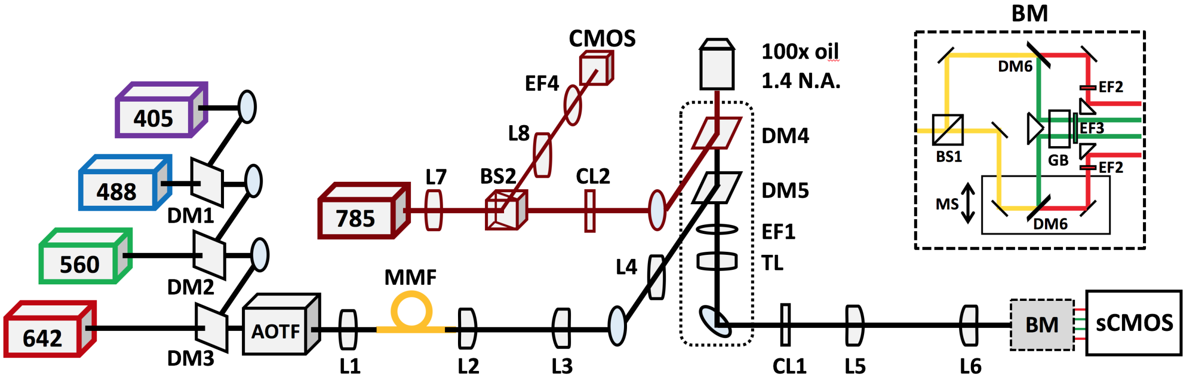


Supplementary Figure 1: Optical setup. L1-8: lenses; DM1-6: dichroic mirrors; BS1-2: beam-splitter cubes; EF1-4: emission filters; AOTF: acousto-optical tunable filter; MMF: multimode fiber; TL: tube lens; CL1-2: cylindrical lenses; BM: biplane module; MS: manual stage; GB: glass block. The details of the biplane module are shown in the top right insert.

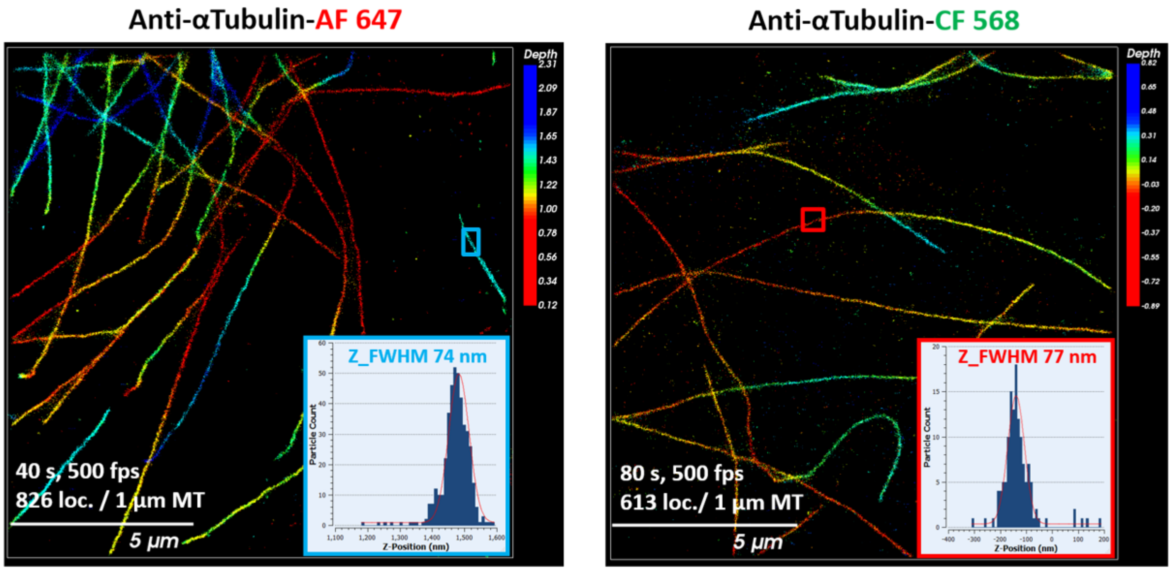


Supplementary Figure 2: SMLM images of microtubules labeled with AF647 or CF568 recorded with our instrument. Insets show axial localization distributions of the blue- or red-boxed regions, respectively. Colorbar units in µm.

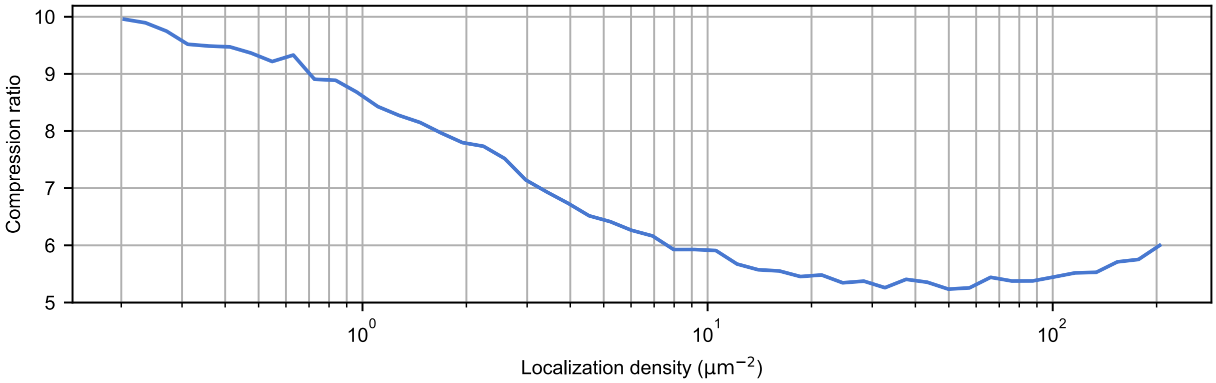


Supplementary Figure 3: As expected, the achievable compression ratio varies with localization density (molecules/$\mu m^2$/frame), ranging from around 5 to around 10 (quantized at $q = \sigma/2$). Note: typical experimental localization densities would be around $1/\mu m^2$/frame.

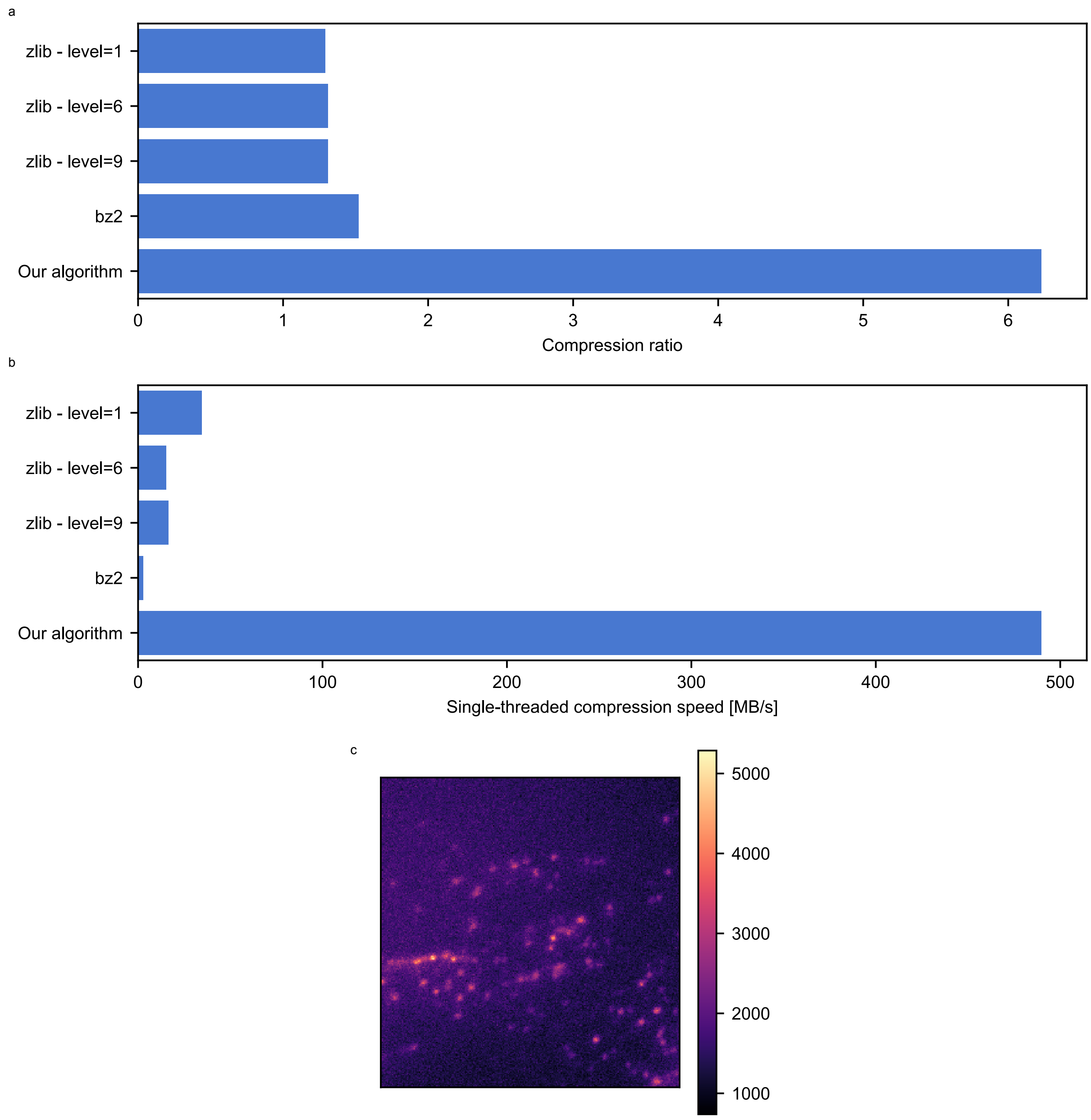


Supplementary Figure 4: (a) Our compression algorithm offers substantially better compression ratios on noisy PALM/STORM image data than general purpose compression algorithms such as gzip or bzip. (b) Through the use of optimized CPU code it is also much faster, such that when used with multi-threaded spooling it easily keeps up with the camera data rate (benchmarks performed on a 2014 Macbook pro, with a 3 GHz Intel processor). (c) Test frame used for benchmarks in (a, b).

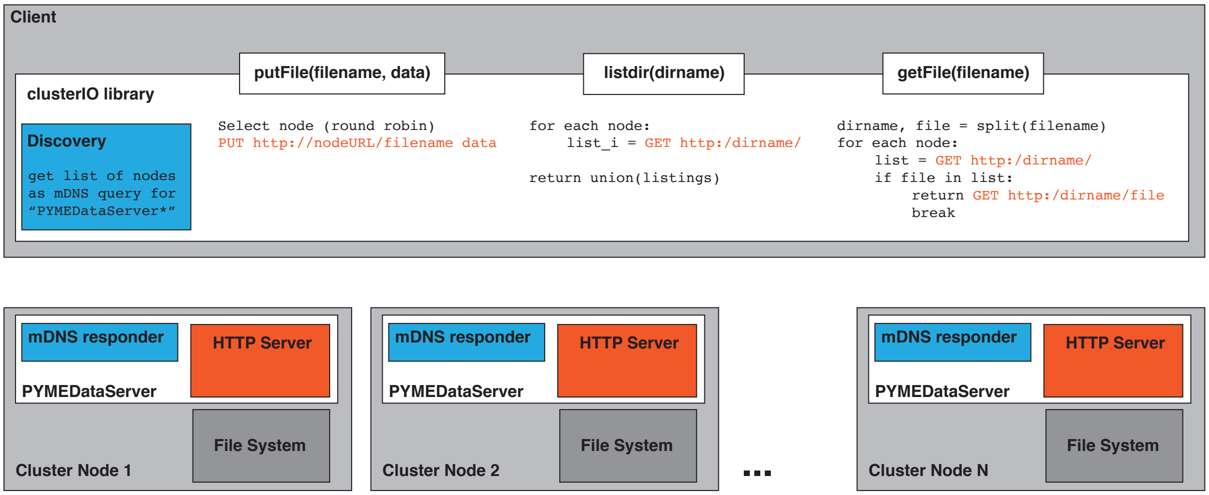


Supplementary Figure 5: High-level schematic of distributed data store with pseudocode for the 3 fundamental operations showing that the vast bulk of filesystem logic lies within the client library, which effectively presents a merged version of the directories on all server nodes. High performance is achieved by using established protocols and atomic file reads/writes. The actual code has a number of optimizations which are not shown, such as the use of persistent HTTP sessions, caching of directory listings, and short-circuiting IO requests for files which are stored on the local machine.

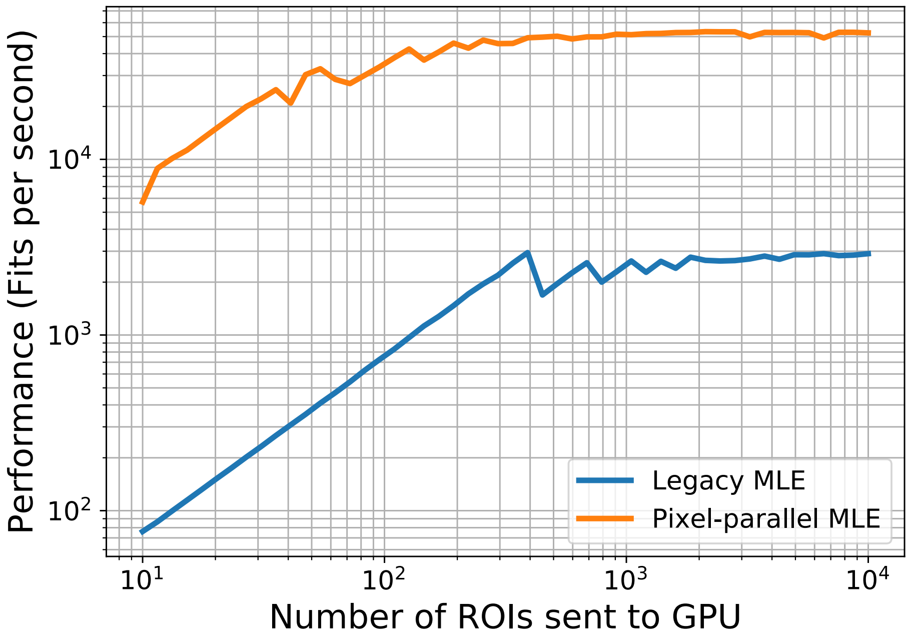


Supplementary Figure 6: Comparison of legacy MLE and pixel-parallel MLE speeds. The legacy implementation of the MLE kernel (blue), is parallel at the level of one-thread per fitting ROI, while the pixel-parallel MLE (orange) is parallel at the level of one-thread per pixel within the fitting ROI. Fitted ROIs were 15 x 15 pixels, and the number of ROIs sent to the GPU for fitting was varied, as shown on the abscissas. Fits were performed on a PNY Quadro M4000 GPU.

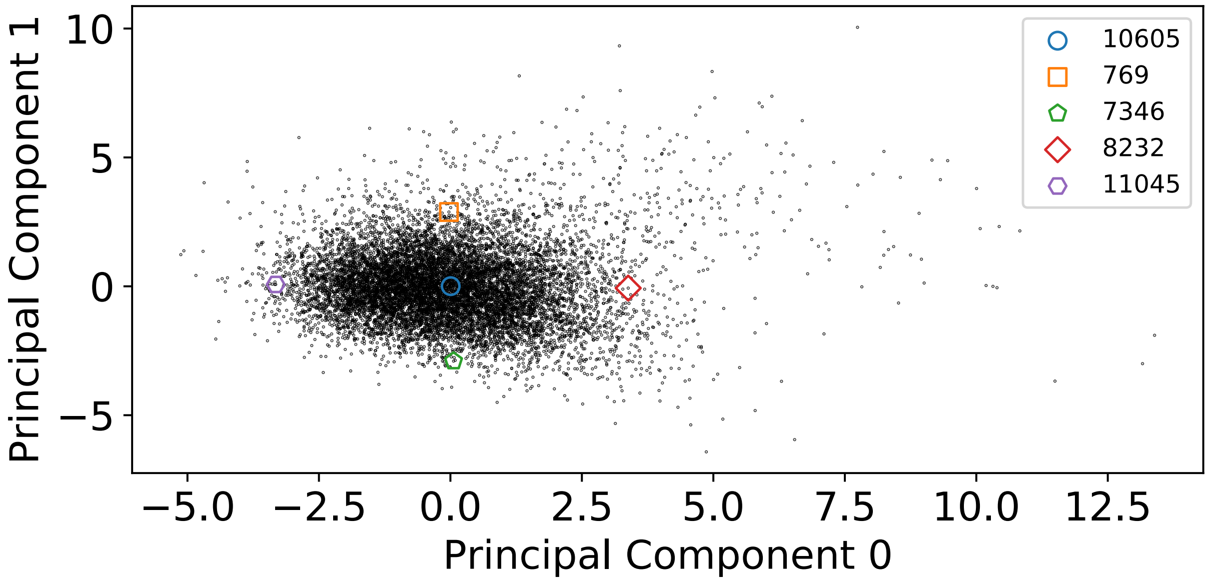


Supplementary Figure 7: Scatter plot of cell morphology principle components from Figure 4c showing the points corresponding to 11,117 nuclei. ROI numbers of cells nearest to the average (PC0, PC1) position and ±2MAD along PC0 and PC1 are given the the legend with colored symbols matching colored outlines in 4c.

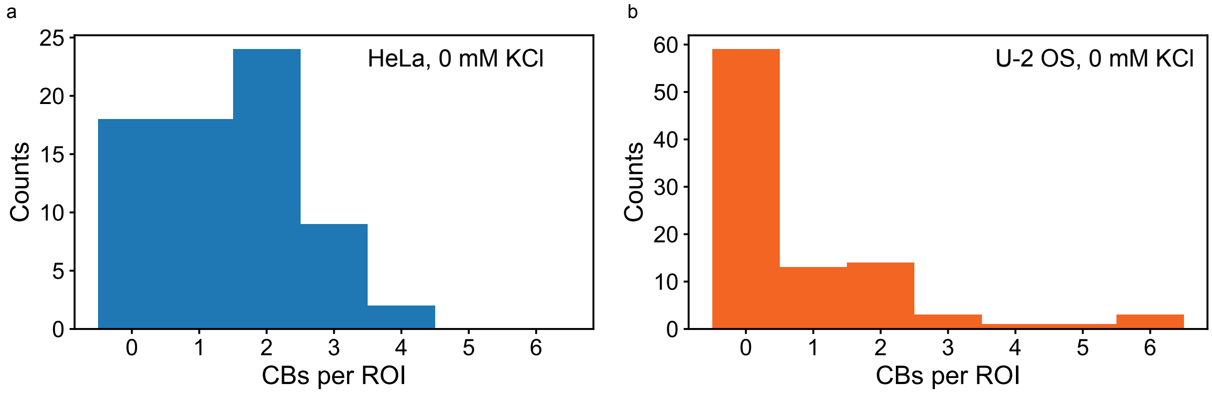


Supplementary Figure 8: Number of Cajal bodies per ROI from the control well (no osmotic shock) datasets in Figure 5. (left) HeLa, (right) U-2 OS.

| Speed [fits/s] | CPU fitting | GPU fitting |
|---|---|---|
| Single Computer | 239 | 1090 |
| 9 Computers | 2071 | 14490 |

Table 1: Localization analysis benchmarks. Localization analysis was timed using either an all-CPU routine (sCMOS least squares, an astigmatic version of [1], PYME's 'AstigGaussFitFR') or all-GPU routine (sCMOS MLE, see Supplementary Note 5), and on a single machine with local data running 10 workers or across a cluster of 9 computers each with 10 workers. Data were the first 10 series of the no-shock HeLa cells in Figure 5. 14,490 fits/s corresponds to 1504 frames/s on these data.

| | Feature | PC0 | PC 1 | Description of quantity being proxied |
|---|---|---|---|---|
| 0 | $\lambda_2^{\text{lamin}}$ | 0.0821 | -0.167 | thickness of the nucleus |
| 1 | $\lambda_0^{\text{lamin}}/\lambda_1^{\text{lamin}}$ | 0.0700 | 0.208 | lateral aspect ratio of the nucleus |
| 2 | $\lambda_0^{\text{NPM1}}/\lambda_1^{\text{NPM1}}$ | 0.0870 | 0.115 | lateral aspect ratio of the nucleophosmin |
| 3 | $\sum_{i=0}^{2}\lambda_i^{\text{NPM1}}/\sum_{i=0}^{2}\lambda_i^{\text{lamin}}$ | 0.465 | 0.396 | relative nucleophosmin to lamin volume |
| 4 | $\lambda_0^{\text{lamin}}+\lambda_1^{\text{lamin}}$ | 0.182 | -2.12 | lateral size of the nucleus |
| 5 | $\sum_{i=0}^{2}\lambda_i^{\text{NPM1}}$ | 0.477 | 0.147 | volume occupied by nucleophosmin |
| 6 | $N^{\text{NPM1}}/N^{\text{lamin}}$ | 0.295 | -0.258 | ratio of nucleophosmin to lamin signal |
| 7 | $r_g^{\text{lamin}}$ | 0.196 | -0.192 | radius of lamin signal |
| 8 | $r_g^{\text{NPM1}}$ | 0.455 | 0.152 | radius of nucleophosmin signal |
| 9 | $N^{\text{NPM1}}/\sum_{i=0}^{2}\lambda_i^{\text{NPM1}}$ | -0.0948 | -0.437 | nucleophosmin density |
| 10 | $\rho_{\in[0,1)}^{\text{NPM1}-\text{NPM1}}$ | 0.121 | -0.413 | nucleophosmin density |
| 11 | $\rho_{\in[9,10)}^{\text{NPM1}-\text{NPM1}}$ | 0.281 | -0.361 | nucleophosmin density |
| 12 | $\rho_{\in[0,1)}^{\text{lamin}-\text{NPM1}}$ | 0.264 | -0.272 | colocalization of lamin and nucleophosmin |

Table 2: The 13 metrics calculated for each lamin B1-nucleophosmin SMLM image. Lamin B1 is denoted 'lamin' and nucleophosmin is abbreviated 'NPM1'. $N^{\text{channel}}$ is the number of localizations in the specified channel, and $\lambda_i^{\text{channel}}$ are the standard deviations along the principal axes (the square root of the $i^{\text{th}}$ covariance eigenvalues) sorted such that $\lambda_0^{\text{channel}}$ is the largest and $\lambda_2^{\text{channel}}$ is the smallest. $r_g^{\text{channel}}$ is the radius of gyration (or root-mean-squared radius). The final 3 metrics refer to specific ranges within the pairwise distance distributions between localizations, where $\rho_{\in[\alpha,\omega)}^{\text{channelA}-\text{channelB}}$ denotes the number of pairwise channelA to channelB distances that lie between $\alpha$ and $\omega$ micrometers).

| | Measure | Description of quantity |
|---|---|---|
| 0 | $r_g$ | radius of gyration |
| 1 | MAD | median absolute deviation |
| 2 | $\frac{\text{Var}\,\sigma_j^2}{\sqrt{3}\langle\sigma_j\rangle}$ | anisotropy. $\sigma_j$ is the standard deviation along principle axis $j \in \{0, 1, 2\}$ |
| 3 | N | Number of localizations |
| 4-6 | $\sigma_j$ | standard deviation along the principle axis $j \in \{0, 1, 3\}$ |

Table 3: Measurements taken of each (candidate) Cajal body which were used in a PCA to select the ROI displayed in Figure 5.

#### Supplementary Notes

Two iterations of the instrument/control platform were used in this work with minor component changes which will be referred to as phase 1 (P1) which was used to collect data in Figures 2, 4, and 2, and phase 2 (P2) which was used for the remaining datasets appearing in Figures 5.

### 1 Automated Biplanar Astigmatism SMLM Microscope

#### 1.1 Instrument

The optical setup is shown schematically in Figure 1, and component abbreviations in the figure are referenced in this section. The setup is custom-built based on a commercial inverted microscope stand (IX71, Olympus) equipped with a 100x/1.4 NA oil immersion super-apochromat objective. The objective is mounted in a piezo actuator (P-726.1CD; Physik Instrumente), which provides an axial travel range of 100 µm. For translating the sample in the $xy$ plane, a motorized sample stage (SCAN IM, Marzhauser) provides a range of 120 × 80 mm.

Fluorescence excitation is provided by four lasers, namely a 405 nm laser (100 mW, CUBE 405-100C, Coherent), a 488 nm laser (500 mW, CUBE 488-500C, Coherent), a 560 nm laser (2 W, 2RU-VFL-P-2000-560-B1R, MPB Communications), and a 642 nm laser (2 W, 2RU-VFL-P-2000-642-B1R, MPB Communications), which are combined into a common beampath by dichroic mirrors (DM 1-3 in Supplementary Figure 1). An acousto-optic tunable filter (AOTFnC-400.650-TN, AA Opto-Electronic) is used as a switch for the lasers, and to regulate their intensities. The laser light is coupled into a multimode fiber (MMF; P1: M43L02, Thorlabs, P2: 04905-1-REVA 100 µm square core 0.16 NA, Armadillo Sia ) by lens L1. At the other end of the fiber, light is coupled out by an f = 4.5 mm lens (L2) and passes through a pair of beam-expansion lenses (L3, f = 300 mm; L4, f = 400 mm). Lenses are positioned such that the fiber tip is imagined into the sample plane of the objective ("critical" illumination). Vibration motors (P1: one or two RadioShack or Jinlong Machinery & Electronics Z4TL2B124064X, P2: 909-100 1.87 G peak-to-peak, Precision Microdrives selected and procured by Cairn Research) were taped to the fiber to remove speckle patterns. Homogeneous illumination is achieved over a circular area of about 30 µm diameter on the sample plane. The fluorescence signal is collected by the objective, and separated from the excitation laser light by a dichroic mirror (DM5, ZT405/488/561/647rpc, Chroma) before passing through a multi-band emission filter (EF1, ZT405/488/561/647m, Chroma).

To expand the axial localization range, a combination of astigmatism [2] and biplane [3] methods are used. Astigmatism is introduced by mounting an f = 1000 mm cylindrical lens (CL1, LJ1516RM-A, Thorlabs) in a holder that can be rotated around the optical axis in the detection beam path. To achieve biplane detection in two color channels simultaneously, we adapted a biplane module (BM, Vutara Inc.) to generate four images of the field of view by first splitting the fluorescence emission into two parts via a beam-splitter cube (BS1, 47-009, Edmund Optics), and then spectrally separating each part using dichroic mirrors (DM6, ZT640rdc-UF1, Chroma). For each color channel, additional emission filters (EF2, ET650lp, Chroma; EF3, ET650sp, Chroma) are used to reduce background. The fluorescence from both color channels on both planes is imaged onto a sCMOS camera (Orca Flash 4.0 v2, Hamamatsu Photonics) by a lens pair (L5, f = 300 mm; L6, f = 175 mm) which corresponds to an effective pixel size of 115 nm in the sample plane. To compensate for the difference in the optical path length (OPL) between the two color channels, a glass block (GB, PS910-Eo2L, custom order from Thorlabs) is inserted in the path with shorter OPL. To finely control the focal shift between the two image planes, the optics for one biplane optical path are mounted on a manual translation stage (MS, M-DS40-x, Newport). All of the optics and optomechanics within the biplane module are of small sizes to fit into the compact design.

A focus-lock system is integrated in the setup to stabilize the axial distance between the sample and the objective during data acquisition. For this purpose, a 785 nm fiber-coupled diode laser (LP785-SF30; Thorlabs) is introduced into the objective at an angle after reflecting off a short-pass dichroic mirror (DM4; FF756-SDi01, Semrock). For this, P1 used L7 and CL2 as shown with focal lengths 16.5 mm, and 500 mm, respectively. P2 instead expanded the beam and did not use a cylindrical lens for the tube lens. The reflection from the interface between coverslip and sample-embedding medium is collected by the objective, separated from the laser light by a beam-splitter cube (BS2; P1: BS017, Thorlabs, P2:), and imaged onto a USB CMOS camera (CMOS; P1: DCC1545M, Thorlabs, P2: UI-3240LE-NIR, IDS) after passing through an

809/81 emission filter (EF4; FF02-809/81-25, Semrock). The position of the reflected light on the camera chip (which depends on the axial position of the sample) is localized used as feedback to adjust the piezo objective actuator (P1: LabVIEW P-loop servo with dead zone, P2: server-connected PYMEAcquire instance with Ziegler–Nichols-tuned PI-loop servo).

#### 1.2 Multi-Channel Lateral Registration and Axial Mapping

##### 1.2.1 Calibration Generation

The two-color biplane module of our microscope takes four images of each FOV by splitting the light into two beam paths focused at different planes and further splitting each of these beams into two color channels. A cylindrical lens in front of the biplane module additionally introduces astigmatism. The four images are projected next to each other onto a single sCMOS camera. To determine the position of the imaged molecules in 3D, the localization algorithm relies on calibrations for (i) registering the four images laterally and (ii) linking the shape of the intensity distribution (on each plane it is found) to the axial position of a molecule. These two calibrations are usually generated just before or after imaging sessions. The functions to apply these calibrations to a set of localizations are written in Python and C.

Calibrations for lateral registrations are generated from reasonably dense bead stacks (using 99 nm diameter fluorescent beads; TetraSpeck, Thermo Fisher Scientific T7279), typically with about 40 axial steps of 25 nm, starting at or below the lower focus and stepping upwards just through the second focal plane, with 3 frames per step. The beads in this series are then localized and localizations from other channels are overlaid onto the 1st channel by subtracting the expected mean lateral offset of that channel based on its position on the camera chip. The localizations are then filtered by $x$ and $y$ localization uncertainties (typically keeping localizations with a maximum Cramer-Rao Lower Bound (CRLB) of 15 nm in $x$ or $y$).

Before pairing localizations from different channels, mean offsets are accounted for using correlations between 2D histogram images rendered from the localizations in each channel. After these shifts are applied, the localizations are clustered (typically 250 - 300 nm lateral search radius) and clusters are kept if they contain localizations from each of the four channels. This allows us to calculate deviations from the expected mean offset with respect to channel 1 for each bead in each of the channels 2 through 4. We fit a shift vector field to these deviations using the following polynomials,

$$\Delta x = a_0 + a_1 x + a_2 y + a_3 x^2 + a_4 y^2 + a_5 xy + a_6 xy^2 + a_7 yx^2 + a_8 x^3, \tag{1}$$

$$\Delta y = b_0 + b_1 x + b_2 y + b_3 x^2 + b_4 y^2 + b_5 xy + b_8 xy^2 + b_7 yx^2 + b_8 y^3 \tag{2}$$

which we have identified as the minimum order required to consistently give a good fit to experimental shift data.

Axial calibrations are generated from bead stacks, typically 25 nm axial steps, 2 to 5 frames per step, of a FOV containing between one and three fluorescent beads (99 nm diameter; TetraSpeck, Thermo Fisher Scientific T7279). Frames from the same z-step are averaged. Beads are manually selected and the bead ROIs in each image are laterally aligned with sub-pixel precision in Fourier space, transformed back to spatial coordinates, and averaged. The 4 (averaged) bead images from each axial step are then fit individually with an elliptic Gaussian function convolved with a $z$-projected bead (modeling the bead as a sphere) to determine $x$ and $y$ PSF widths as a function of $z$. To determine the useful axial range of this calibration for each of the 4 (color or plane) channels we fit a cubic smoothing spline to the difference in lateral widths ($\sigma_x - \sigma_y$), and identify the range over which this difference is monotonic. Once the valid range is established we fit cubic smoothing splines independently to $\sigma_x(z)$ and $\sigma_y(z)$ for each of the 4 channels. For all fitted splines, the knots are typically placed at 50 nm intervals.

##### 1.2.2 Registration, Linking, and Axial Mapping

The first step of the process of registration, linking, and axial mapping is adding the lateral shifts from each (plane and color) channel to localizations from that channel. After these shifts are applied, localizations which where individually detected in both planes of the same color channel are linked, as are localizations in subsequent frames if they are within a distance of twice the localization uncertainty. Gaps of one frame where a molecule was not detected are accepted in the merging procedure, though if gaps are two or more frames in duration, subsequent localizations are considered to be from a different molecule. The calibration splines for $\sigma_x(z)$ and $\sigma_y(z)$ for each plane are evaluated at 1 nm intervals over the entire valid range to give calibration curves, $\zeta_x$ and $\zeta_y$, to which the fitted values of $\sigma_x$ and $\sigma_y$ can be compared.

For each molecule and each detection plane, the squared difference between the fitted $\sigma_x$ and $\sigma_y$ values and the calibration curves $\zeta_x$, $\zeta_y$ are calculated giving the following 4 error curves:

$$E_{\sigma_{x_0}}(z) = [\sigma_{x_0} - \zeta_{x_0}]^2 \tag{3}$$

$$E_{\sigma_{x_1}}(z) = [\sigma_{x_1} - \zeta_{x_1}]^2 \tag{4}$$

$$E_{\sigma_{y_0}}(z) = [\sigma_{y_0} - \zeta_{y_0}]^2 \tag{5}$$

$$E_{\sigma_{y_1}}(z) = [\sigma_{y_1} - \zeta_{y_1}]^2 \tag{6}$$

Where $\sigma_{x_0}$ is the fitted $x$ width in the first biplane plane and $\sigma_{x_1}$ is the width in the second biplane plane. These 4 error curves are combined by taking a weighted sum using the uncertainties, $\Delta$, in our estimation of that molecule's $\sigma_{x,y}$ (CRLBs of $\sigma_x$ and $\sigma_y$ from the MLE) to give:

$$E(z) = \frac{1}{\Delta^2_{\sigma_{x_0}}} E_{\sigma_{x_0}}(z) + \frac{1}{\Delta^2_{\sigma_{x_1}}} E_{\sigma_{x_1}}(z) + \frac{1}{\Delta^2_{\sigma_{y_0}}} E_{\sigma_{y_0}}(z) + \frac{1}{\Delta^2_{\sigma_{y_1}}} E_{\sigma_{y_1}}(z) \tag{7}$$

The axial position with the minimum summed and weighted error is assigned to the molecule.

### 2 Compression Testing

We benchmarked our lossy compression algorithm on simulated single molecules. 5,000 ROIs containing a single Gaussian emitter with peak amplitude 225 photoelectrons, constant background of 4.5 photoelectrons, center jittered by a 1 pixel sigma normal distribution in $x$ and $y$, and sigma of 1 pixel were simulated on 15 x 15 pixel ROIs subjected to Poisson noise and fitted, noting the error in the fitted x position from the ground truth. Each ROI was then quantized and re-fitted, at various quantization levels. The mean absolute error in fitted x position at each quantization level was calculated and normalized by the mean absolute error of the unquantized fitted $x$ positions.

To benchmark achieved compression ratios frames of 256 x 256 pixels were simulated with Gaussian emitters of sigma 2 pixels, quantized, and compressed using Huffman coding, with the averaged compression ratio from 10 frames for each quantization reported.

An additional speed and compression benchmark is provided in supplemental figure 4 and were run on a single frame from an EMCCD recording (shown in the figure).

### 3 PYME Cluster

#### 3.1 Cluster Hardware

The cluster used for data storage and analysis consists of 10 nodes built from inexpensive commodity PC hardware in a single 42U computer rack. Each node has a single intel Core i7-5820K CPU, 32 GB of DDR4 SDRAM, a CUDA-capable GPU (PNY NVIDIA Quadro M4000), an SSD boot drive, and five 3 TB hard drives configured in RAID5 for data storage. The nodes are connected to a switch (NETGEAR S3300-28X) with a 10 GbE connection to the instrument computer. Software on the instrument computer ensures that the data flow (up to 800 MB/s) from the camera is distributed evenly amongst the individual nodes (see Supplementary Note 3.2), allowing cheaper 1 GbE connections to be used within the cluster. The cluster was built in early 2016, and during subsequent years, 2 of the nodes were replaced with nodes that are nearly identical with the exception of GPU; they are equipped with NVIDIA GeForce 1080Ti, with more recent experiments conducted using this modified cluster (8x M4000, 2x1080Ti). We note that this cluster was originally built in early 2016, and while functional for high-throughput imaging, advances in hardware would suggest different selections if built today; a higher performance cluster could now be built for roughly \$20,000 - \$25,000 (USD). Our software will run on a wide range of cluster configurations, but an increase in network bandwidth within the cluster might be needed if the number of nodes were substantially decreased. The authors have since helped with setting up several other clusters with more updated equipment, and are open to being contacted for hardware advice.

##### 3.2 Distributed Data Storage

To allow uninterrupted spooling and storage of large volumes of data at the full camera frame rate we have developed a distributed file system specifically optimized for streaming image data. It is similar in concept to existing systems such as Apache Hadoop which shard data across nodes, but preserves frame granularity so that an entire image frame (or sequence of frames) will be stored on a single node permitting data-local image analysis. There are two additional design differences both aimed at achieving high write speeds by reducing the amount of coordination and locking that is needed between nodes: we employ a write-once idiom where data that has been written cannot be modified, and we only support atomic reads and writes of a whole file (i.e. no random access within files).

Our distributed file system is implemented as a combination of a Python library ('clusterIO') which acts as a client, and a series of HTTP servers - one on each node of the cluster - which serve the files stored on that node. In addition to the normal HTTP server functions, these servers implement the PUT verb, and a few custom REST endpoints to facilitate indexing and duplication operations. Atomic operations remove the need for locking and a master node and allow almost all of the filesystem logic to reside in the client library which effectively presents the different files stored across different nodes as one unified file system. The servers advertise themselves using the mDNS/zeroconf protocol and are automatically discovered by the clusterIO client library meaning that file system setup and configuration is as simple as running a copy of the server process on each node of the cluster. Enabling duplication makes the filesystem robust to node failure. When used during the spooling of raw data, the clusterIO library automatically distributes writes amongst servers such that a roughly equal data volume is saved to each server.

In addition to the low level access permitted by clusterIO, we have developed a web application interface to the cluster storage and analysis algorithms and a WebDAV backend which permits the cluster file system to be mounted as a shared drive.

##### 3.3 Data-Local Distributed Analysis

In order to minimize network IO within the cluster, which would otherwise limit analysis speed, raw data frames should be analyzed on the node on which they are saved. To allow this to happen, we have developed a framework for assigning analysis tasks at high speed. Due to the high rate (>800 new frames to be analyzed per second) and high volume ($\sim$70 million tasks in a 24 h period), this framework is non-trivial. Our solution consists of multiple elements: a master server ('PYMERuleServer') maintains a list of rules which describe analysis tasks which need to be performed. A server process ('nodeserver') running on each node matches these rules to files on the cluster, assigns a cost (low if the file is local, higher if it is on another node) and asks the rule server if they can perform the analysis, bidding preferentially on files that they have locally. If no other node has made a better bid on that file, they get permission to run the analysis. From the nodeserver, tasks are further distributed to a number of local worker processes, one for each core of the node. The use of multiple worker processes (each with their own Python global interpreter lock) both allows maximum performance for the parts of the analysis performed on the CPU and allows memory transfers to the GPU to be interleaved with GPU-based processing. Each worker then saves its analysis results directly to the cluster and reports the task as complete. The system is robust against worker failure, with incomplete tasks being re-queued after a timeout, and can be used to schedule both primary localization tasks and subsequent quantitative analysis, facilitated by a flexible 'recipe'-based definition of analysis pipelines. Like the distributed data storage, the analysis framework is self-configuring using mDNS and its setup is as simple as running worker processes on all the nodes and a master on one node.

##### 3.4 Setting up the computer cluster

Setting up the software on the cluster is straightforward for someone with Linux and Python experience. Set-up instructions can be found at `http://python-microscopy.org/doc/cluster/cluster_install.html`

### 4 Integrated Acquisition and Analysis Workflows

Automation of complex PYMEAcquire acquisitions is achieved using a combination of 3 constructs: acquisition *protocols* which encode how to acquire a particular type of image series, a queue which schedules the acquisition and analysis of individual image series, and an interface which enables analysis tasks to add to the queue.

##### Acquisition protocols

The common steps and settings needed for a given type of image acquisition are formatted as a *'protocol'* - a text-based listing of operations to be performed prior to the start of acquisition (setup), on specified frame numbers during acquisition, and after acquisition is complete (cleanup). Protocols can be used to change laser and camera settings, as well as configuring z-stepping and similar operations. In this work, protocols were either for an SMLM acquisition, or for tiling an overview image of a coverslip.

##### Action queue

Individual series acquisitions, series specific changes in state (i.e. moving the stage to the position of a cell or region of interest), and optionally, the triggering of analysis on the cluster are scheduled using a priority queue in the microscope control software. This action queue exposes a REST endpoint, allowing analysis tasks running on the cluster to add new acquisition tasks. The use of a priority queue allows higher priority tasks to push to the front of the queue, with execution of the original tasks resuming on completion of the high priority task. When imaging multi-well samples in this paper we initially populated the queue with tasks to complete a tiled overview of each well and to trigger ROI detection on the cluster. We then used task priority to insert ROI imaging tasks based on these detections so that they executed with a higher priority than the remaining overview tasks, minimizing the number of large-distance movements of the well plate and the associated risk of immersion oil loss. Actions which must be executed as a group (e.g. a move and an acquisition) can be chained so that they appear as one entry in the queue, preventing them from being interrupted by a higher priority task.

##### Analysis rules and recipes

Data analysis on the cluster is facilitated by analysis *rules* which describe what data to process (see 5) and *recipes* (see 3.3)) which describe what processing to perform. Like acquisition tasks, analysis rules can be chained - for example to perform localization analysis, then generate voxel-based images from the localizations, then perform quantification.

##### Integrated workflows

Integrated workflows, where automatic analysis feeds back into new acquisition tasks, are enabled by queueing new acquisition actions from within a recipe executing on the cluster, based on results of analysis performed within that recipe. The use of this mechanism for multi-well imaging (as in Figure 5 and described above) is one example of such an integrated workflow, but the architecture is very flexible - it could, for example, also be used in live-cell SMLM to trigger high-resolution imaging of rare events in living cells which are monitored using low-resolution (and low-dose) overview imaging.

### 5 GPU-Accelerated sCMOS-Specific Localization

The sCMOS-specific localization routine used in this work is based on the algorithm described in [4], but with substantial speed improvements and the added capability to use an estimated pixel-specific background to improve candidate molecule detection and MLE localization in moderate-density image series.

Although our previous sCMOS-specific localization algorithm[4] was already written to use the GPU, simply distributing analysis across ten computers was not sufficient to allow real-time localization and a boost to the performance on a single machine was also necessary. This was accomplished by changing the architecture of the existing GPU code such that it was parallelized so that each GPU thread processed one pixel rather than a whole ROI and by moving candidate detection to the GPU.

In the previous implementation of this algorithm [4], candidate molecule detection is performed on the CPU in MATLAB, with the MLE performed on the GPU using CUDA. While the candidate detection steps are not computationally complex, they were taking a similar amount of time as the GPU based fitting. We accelerated the sCMOS-specific candidate molecule detection steps by writing GPU kernels for them, called from Python through PyCUDA [5]. Porting all candidate detection steps to the GPU resulted in over an order of magnitude improvement in candidate detection speed, additionally reducing expensive CPU to GPU memory transfers.

By using the CUDA Multi-Process Service (MPS), which allows multiple CPU threads to manage localization operations on separate frames while sharing the GPU resources, and performing computations in multiple processes simultaneously we were additionally able to hide much of the latency involved in CPU-GPU data transfer as one process could by uploading data while another processes fits were running.

In order to more readily find candidate molecules, and fit them on top of a non-uniform background as is often the case early during high-throughput SMLM series (we forgo a dedicated pre-bleaching period), we employ robust computational background estimation using a sliding-window percentile. We sort the intensity values at each pixel over the preceding 32 frames and take the $25^{\text{th}}$ percentile to be the background. This background is then subtracted during the candidate molecule detection. During the MLE fitting, the background estimate is not subtracted, but instead added onto the model when calculating the residuals at each iteration of the fit - this enables us to remove the influence of structure background from the fitting without altering the intensity values and therefore the noise-model of our fit.

#### 5.1 CUDA implementation details

Our kernel architectures should perform well on all modern NVIDIA GPUs and are parallel at the level of 1 GPU thread per $(x, y)$ or $(x, y, t)$ pixel. While these kernels are operating-system independent, the performance is maximized when using CUDA Multi-Process Service (MPS), which is currently limited to Linux operating systems.

##### 5.1.1 Per-Pixel Background Estimation

To find the $25^{\text{th}}$ percentile of our background window we implemented a simple sort algorithm in CUDA, parallelized at the level of one thread per pixel and time point. It is similar to an insertion sort, but exhaustively searches to determine the optimal position for each value before making insertions after all optimal positions have been found. Despite an $O(n^2)$ average-case complexity the algorithm performs well for small buffer lengths due to the minimal divergence between thread paths. Our kernel achieves 100% theoretical occupancy for buffer lengths of integer multiples of 32 frames, but additionally supports arbitrary buffer lengths.

##### 5.1.2 Candidate Molecule Detection

Following the algorithm in Huang, et al. [4], we perform several weighted convolutions with uniform kernels to effectively bandpass-filter each frame, resulting in a smoothed image, $S$, where the $j^{\text{th}}$ pixel is given by

$$S_j = \frac{\sum\limits_{i \in C_0} \frac{D_i - o_i - b_i}{g_i \text{var}_i}}{\sum\limits_{i \in C_0} \text{var}_i^{-1}} - \frac{\sum\limits_{i \in C_1} \frac{D_i - o_i - b_i}{g_i \text{var}_i}}{\sum\limits_{i \in C_1} \text{var}_i^{-1}}, \tag{8}$$

where $D$ is the intensity counts in a single frame, $o$ is the camera offset map, $b$ is the background estimate for the frame, $g$ is the gain map in units of [ADU / $e^-$], and var is the variance map. $C_0$ and $C_1$ denote square regions around the $j^{\text{th}}$ pixel. Typically $C_0$ is around twice the size of $C_1$. The denominators in equation 8 are computed once per GPU context upon initialization. The convolutions are performed in real space in a row-then-column manner as the uniform filter is separable, with the row or column loaded into shared memory.

A maximum filter is then applied to the smoothed image [4]. Maximum (and minimum) filters are also separable, so this function borrows the same code structure of the former row- and column-convolutions.

A per-pixel threshold based on the estimated SNR for each $(x, y, t)$ pixel is applied to the smoothed image [1]. No shared memory is required, though each row of pixels is given a block of threads such that there is one thread per (x, y) pixel. Each pixel, $j$ is assumed to be a candidate molecule position if at that location the smoothed image, $S_j$, is equal to the maximum filtered image, $\max_{i \in C_2} S_j$, and this value is larger the SNR-based threshold at pixel $j$, where $C_2$ is a square region with dimensions similar to the fitted ROI ($C_2$ is a 16 x 16 or 17 x 17 pixel square in this work, as our fitted ROIs are 16 x 16 pixels). To avoid threads writing over each other in the output array, the index of the output array is handled using atomic functions.

###### 5.1.3 Gaussian MLE

The legacy implementation of the MLE kernel is parallel at the level of one-thread per fitting ROI (16 x 16 pixels in this work) and is performance-limited by shared memory requirements. We restructured the MLE kernel to be parallel at one-thread per pixel, and stored pixel intensities in per-thread memory rather than shared memory, achieving a substantial speed improvement (see Supplementary Figure 6).

### 6 Overviews and ROI Selection

For automated imaging, the sample was allowed to settle with the focus servo turned on before preview scanning was initialized. The preview scan is performed using the same objective as our SMLM imaging, moving the stage in discrete steps and inserting images with no overlap into their predetermined positions in the overview image (P1, used in Figure 4) or with intentional overlapping and mean-based stitching using ex post facto positions associated with each ROI (P2, used in Figure 5).

#### 6.1 No-Overlap Overviews

No overlap P1 overviews were acquired in LabVIEW and used steps of ~20 µm between acquisition of images with 4 x 4 pixel binning, resulting in 48 x 48 pixel images. These individual preview FOVs were acquired with 2 ms exposure times and either ~600 W/cm$^2$ of 560 nm light exciting the lamin stain for the lamin B1 and NPM1 sample in Figure 4, or ~90 W/cm$^2$ of 405 nm light exciting Hoechst 33342 for the lamin A/C and LAD pool 2 sample, and the Chr 22 TAD and LAD pool 1 sample. The preview FOVs, with no intentional overlap, were inserted directly into a mosaic coverslip overview image. Automatic selection of FOVs for SMLM imaging was then performed by generating a binary mask from the mosaic overview image using intensity thresholding, and then rejecting regions which did not meet empirical size limits. These limits were chosen to detect well-separated interphase nuclei. The preview scan took approximately 52 minutes for the 300 x 300 FOV (6 x 6 mm) scan used for the Lamin B1 and NPM1 imaging in Figure 4.

#### 6.2 Flexible Overviews

Coverslip scanning in phase 2 is substantially more flexible. Tile overview series can be spooled from PYMEAcquire to the computer cluster, tiling either square overviews or circular overviews spiralling radially outward, both with overlapping ROIs acquired using a software trigger after stage movement. For each frame, the coordinates of the stage is logged and used to build the base layer (highest zoom) of a multi-level image pyramid using a PYME recipe. These multilevel pyramids can be viewed using the PYME tileviewer application for manual ROI selection by a user, or used in a PYME recipe for automated ROI selection.

In this work, phase 2 overviews were used to detect nuclei using a recipe run automatically on the cluster upon overview series acquisition. The overview series were first converted to multi-level image pyramids and then a de-magnified layer (typically binned 4x to 460 nm pixels) was processed with a deep neural network to segment nuclei. To this end, the pytorch Mask R-CNN [6] implementation was trained with the BBBC038v1 dataset, available from the Broad Bioimage Benchmark Collection [7]. The model was initiated with weights which had been pre-trained on the COCO dataset [8] and reconfigured to separate between 2 classes: nuclei and noise. It was trained on the data provided in the first stage of the Data Science Bowl 2018 for 50 epochs and a copy was saved after each training epoch. The copy with optimal bounding box precision and recall was used for segmenting and locating nuclei in overview images. The training script is based on the pytorch Object Detection Finetuning tutorial [9] and adapted code from reference scripts (references/detection) in the pytorch vision repository [10]. Though the model was trained with pytorch 1.7.1, inference was done with pytorch 1.1.0 due to dependency issues.

### 7 Sample Preparation

#### 7.1 Cell Culture

U-2 OS (ATCC HTB-96) cells were grown in McCoy's 5A Medium (30-2007, ATCC) supplemented with 10% fetal bovine serum (FBS; 1500-500, Seradigm) at 37 °C with 5% $CO_2$. IMR-90 (ATCC CCL-186) were grown in EMEM (30-2003, ATCC) supplemented with 10% FBS and PenStrep (15140-122, Gibco) at 37 °C with 5% $CO_2$. COS-7 (ATCC CRL-1651) cells were grown in DMEM (21063-029, Gibco) supplemented with 10% FBS (10438-026, Gibco), and sodium pyruvate (11360-070, Gibco) at 37 °C with 5% $CO_2$. HeLa cells were grown in DMEM (21063-029, Gibco) supplemented with supplemented with 10% FBS (10438-026, Gibco) and PenStrep (15140-122, Gibco) at 37 °C with 5% $CO_2$.

##### 7.1.1 Microtubule and ER Immunofluorescence Sample Preparation

A transient transfection of mEmerald-Sec61$\beta$ was used to label ER membranes (mEmerald-Sec61-C-18; a gift from Michael Davidson, Addgene plasmid #54249; http://n2t.net/addgene:54249; RRID:Addgene_54249) COS-7 cells (ATCC batch #63624240) were electroporated with mEmerald-Sec61$\beta$ then seeded onto a 40 mm round coverslip and grown overnight. Prior to seeding with cells, the coverslip was sonicated for 15 minutes in KOH, rinsed 3 times with Milli-Q water, sterilized with 100% ethanol, treated with poly-L-lysine (P4707, Sigma-Aldrich) for 10 minutes, and then rinsed 3 times with PBS. The cells were fixed in 3% paraformaldehyde (PFA; 15710, Electron Microscopy Science) + 0.1% glutaraldehyde (GA; 16019, Electron Microscopy Science) diluted in phosphate buffered saline (PBS) for 15 minutes and then rinsed 3 times with PBS. The cells were permeabilized (PBS + 0.05% IGEPAL CA-630 + 0.05% Triton X-100 + 0.1% bovine serum albumin (BSA, 001-000-162, Jackson ImmunoResearch)) for 3 minutes and then rinsed 3 times with PBS. Following permeabilization, the cells were incubated in blocking buffer (PBS + 0.05% IGEPAL CA-630 + 0.05% Triton X-100 + 5% normal goat serum (005-000-121,Jackson ImmunoResearch)) for 1 hour. The primary antibodies mouse anti-$\alpha$-tubulin (T5168, Sigma-Aldrich) and rabbit anti-GFP (A-11122, Invitrogen) were incubated on samples overnight at 4 °C, at 1:1000 and 1:500, respectively. Cells were then washed 3 times for 5 minutes each with wash buffer (PBS + 0.05% IGEPAL CA-630 + 0.05% Triton X-100+ 0.2% BSA). Samples were incubated with secondary antibodies, anti-mouse labeled with CF568 (20109, Biotium) and anti-rabbit labeled with AF647 (A21245, Invitrogen), for 1 hour at room temperature at 1:1000 and then washed 3 times for 5 minutes each with wash buffer. Finally the samples were post-fixed with 3% PFA + 0.1% GA for 10 minutes and then rinsed 3 times with PBS.

#### 7.2 Mitochondria and Nucleoid Immunofluorescence Sample Preparation

For mitochondria and mitochondrial nucleoid labeling, U-2 OS cells were grown and seeded as described above and then fixed for 1 h with 4% PFA. Then the samples were sequentially labeled starting with a mouse anti-dsDNA antibody (ab27156, Abcam) incubated overnight at 4 °C and goat anti-mouse CF568ST secondary antibody (20800, Biotium) for 1 h at room temperature followed by rabbit anti-TOM20 antibody (ab78547, Abcam) incubated overnight at 4 °C with goat anti-rabbit AF647 secondary antibody (A21245, Invitrogen) for 1 h at room temperature. Both primaries and secondaries were used at a dilution of 1:1000. The sample was post-fixed with 3% paraformaldehyde (PFA; 15710, Electron Microscopy Science) + 0.1% glutaraldehyde (GA; 16019, Electron Microscopy Science) diluted in phosphate buffered saline (PBS) for 10 minutes and then rinsed 3 times with PBS.

#### 7.3 LAD FISH and Lamin Immunofluorescence Sample Preparation

To fluorescently label lamina-associated domains (LADs) with FISH probes, we first downloaded LAD coordinates from three previous publications [11–13] and chose two common regions that were identified as LADs by all three studies. The chosen LAD regions are both between 300 and 310 kb in genomic lengths, and are both within single topologically associating domains (TADs) based on previously reported TAD coordinates [14]. The genomic coordinates of the chosen LADs are: Chr5:115508197-115813276 (LAD1), and Chr13: 24405079-24709084 (LAD2) (all coordinates are based on genome assembly hg18.) We then designed 3725 and 2564 oligonucleotide probes (see Supplementary Extended Table 1 and 2) that targeted the two LADs, respectively, and synthesized the probes. The probe design and synthesis procedures were similar to those in a previous report [15] with only minor modifications. Briefly, we designed the template probe sequences using a custom computation pipeline [15] and generated complex pools of probes using array-based oligo library synthesis [16–18] followed by a high-yield enzymatic amplification approach [19, 20]. We first designed template probe libraries for the synthesis of our probe sets. Each oligo in our template probe libraries contained: i) a 20-nucleotide (nt) forward priming region for library amplification, ii) a 30 nt targeting region for in situ hybridization to the target chromosomal DNA sequence, iii) a 10 nt linker region, and iv) a 20 nt reverse priming region for library amplification. The sequences of the 20 nt forward and reverse priming regions were chosen from random sequences [20] with constraints to ensure their PCR priming performance, and further blasted against the human genome with BLAST+ [21] to ensure their lack of significant homology to the genome. The 10 nt linker sequences were cropped from other 20 nt sequences generated the same way as above. The 30 nt targeting regions were generated with the software OligoArray2.1 [22] and BLAST+ [21] as described previously [15]. The designed template probe libraries were ordered from CustomArray, Inc as oligo pools. We then synthesized the FISH probe sets from the template libraries following a previously published protocol that involved limited-cycle PCR, in vitro transcription, reverse transcription, alkaline hydrolysis and purification [15, 20]. An Alexa Fluor 647 dye molecule was conjugated to each FISH probe through the reverse transcription primer [15].

IMR-90 cells (ATCC CCL-186) were grown in EMEM (30-2003, ATCC) supplemented with 10% FBS and PenStrep (15140-122, Gibco) at 37 °C with 5% $CO_2$. IMR-90 cells were plated on 40 mm diameter, #1.5 coverslips (Bioptechs, 0420-0323-2), and further cultured for 3 to 5 days to reach a confluency of ∼75%. All following procedures were performed at room temperature unless stated otherwise. The cells were fixed in cytoskeletal fixation buffer (2.5% formaldehyde, 10 mM MES, 138 mM KCl, 3 mM $MgCl_2$, 2 mM EGTA, 0.1 g/mL sucrose in DPBS) at room temperature for 25 min, washed twice with DPBS for 2 min each, permeabilized and blocked with blocking buffer (3% BSA, 0.2% v/v Triton X-100 in DPBS) for 30 min with gentle shaking, and incubated with lamin A/C primary antibody (Cell Signaling Technology, 4C11) in antibody dilution buffer (1% BSA, 0.2% v/v Triton X-100 in DPBS) at a concentration of 1:200 at 4 °C overnight with gentle shaking. The cells were then washed with wash buffer (0.05% v/v Triton X-100 in DPBS) for three times at room temperature and incubated with biotin-labeled secondary antibody (Jackson ImmunoResearch Laboratories, 115-065-003) in antibody dilution buffer (1% BSA, 0.2% v/v Triton X-100 in DPBS) at a concentration of 1:1000 for 1 h with gentle shaking, washed with wash buffer (0.05% v/v Triton X-100 in DPBS) three times, post-fixed with 4% formaldehyde for 10 min and washed with wash buffer (0.05% v/v Triton X-100 in DPBS) three times. Next the cells were treated with 0.1 M HCl for 5 min, washed twice with DPBS, digested with 0.1 mg/mL RNAse A (AmericanBio, AB12023-00100) in DPBS for 45 min at 37 °C, washed twice in 2xSSC, and incubated for 30 min in pre-hybridization buffer (2xSSC, 50% formamide, and 0.1% v/v Tween 20). For probe hybridization, 25 µL of hybridization buffer, containing 2xSSC, 50% formamide, 20% dextran sulfate, and a LAD probe set at 4 µM concentration were dropped on top of a glass slide. The coverslip with cells was flipped and placed on top of the slide. The assembly was placed on top of an 80 °C heat block for 3 min, and then incubated for 15-18 h at 37 °C in a humid chamber. The cells were then washed twice with 0.1% v/v Tween 20 in 2xSSC at 60 °C for 15 min each, and once more for 15 min. We then switched the sample buffer to DPBS and incubated it with CF568-conjugated streptavidin (Biotium, 29035) in antibody dilution buffer (1% BSA, 0.2% v/v Triton X-100 in DPBS) at a concentration of 1:1000 for 1 h with gentle shaking. The sample was then washed with DPBS for 3 times, each for 5 min. The sample was additionally incubated with Hoechst 33342 at a concentration of 0.4 µg/mL in DPBS for 5 min and washed twice with DPBS. We then switched buffer to 2xSSC and prepared the sample for STORM imaging.

#### 7.4 Combined TAD and LAD FISH Sample Preparation

The design and synthesis of the LAD probes are the same as described above. FISH probes targeting Chr22 were designed and synthesized as previously reported [15], except that the reverse transcription primers for Chr22 probes were conjugated with a 5' biotin and as a result all Chr22 FISH probes were conjugated with biotin (see Supplementary Extended Table 3 and 4).

The two-color FISH samples were also prepared with a protocol similar to the one described previously [15]. IMR-90 cells (ATCC CCL-186) were grown in EMEM (30-2003, ATCC) supplemented with 10% FBS and PenStrep (15140-122, Gibco) at 37 °C with 5% $CO_2$. IMR-90 cells were plated on 40 mm diameter, #1.5 coverslips (Bioptechs, 0420-0323-2), and further cultured for 3 to 5 days to reach a confluency of ~75%. All following procedures were performed at room temperature unless stated otherwise. The cells were fixed for 10 min with 4% formaldehyde in Dulbecco's phosphate-buffered saline (DPBS), washed twice with DPBS, treated for 10 min with freshly made 1 mg/mL sodium borohydride (Sigma, 480886) solution, washed twice with DPBS, permeabilized for 10 min with 0.5% v/v Triton X-100 (Sigma, T8787) in DPBS, washed twice with DPBS, treated with 0.1 M HCl for 5 min, washed twice with DPBS, and digested with 0.1 mg/mL RNAse A (AmericanBio, AB12023-00100) in DPBS for 45 min at 37 °C. The cells were then washed twice in 2x saline-sodium citrate buffer (SSC) (Ambion, AM9763) and incubated for 30 min in pre-hybridization buffer (2xSSC, 50% formamide (Ambion, AM9342) and 0.1% v/v Tween 20 (Fisher, BP337)). For probe hybridization, 25 µL of hybridization buffer, containing 2xSSC, 50% formamide, 20% dextran sulfate (Sigma, D8906-50G), and probe sets targeting both LAD1 and Chr22 at 4 µM for each set were dropped on top of a glass slide. The coverslip with cells was flipped and placed on top of the slide so that the cells were in contact with the hybridization buffer. The slide-coverslip assembly was placed on top of an 80 °C heat block for 3 min, and then incubated for 15-18 h at 37 °C in a humid chamber to allow hybridization of the FISH probes to their genomic targets. The cells were then washed twice with 0.1% v/v Tween 20 in 2xSSC at 60 °C for 15 min each, and once more at room temperature for 15 min. We then switched the sample buffer to DPBS and incubated the cells with blocking buffer (3% BSA, 0.2% v/v Triton X-100 in DPBS) at room temperature for 30 min. The cells were then incubated with mouse anti-biotin primary antibody (Jackson ImmunoResearch Laboratories, 200-002-211) in antibody dilution buffer (1% BSA, 0.2% v/v Triton X-100 in DPBS) at a concentration of 1:1000 at 4 °C overnight with gentle shaking and washed with wash buffer (0.05% v/v Triton X-100 in DPBS) for three times at room temperature. We then incubated the sample with CF568-conjugated secondary antibody (Biotium, 20109) at a concentration of 1:1000 in antibody dilution buffer (1% BSA, 0.2% v/v Triton X-100 in DPBS) for 1 h with gentle shaking. The sample was then washed with wash buffer (0.05% v/v Triton X-100 in DPBS) 3 times. The sample was additionally incubated with Hoechst 33342 at a concentration of 0.4 µg/mL in DPBS for 5 min and washed with DPBS twice. We then switched buffer to 2xSSC and prepared the sample for STORM imaging.

#### 7.5 Lamin and Nucleolus Immunofluorescence Sample Preparation

U-2 OS cells were seeded onto ozone-cleaned 40 mm round coverslips and grown overnight. For lamin and nucleolus labeling, U-2 OS cells were fixed with 4% PFA for 15 minutes, rinsed 3 times with PBS, permeabilized for 3 minutes, rinsed 3 more times, and then incubated with blocking buffer (as above) for 1 hour. The sample was sequentially labeled starting with mouse anti-nucleophosmin antibody (NB600-1030, Novus Biologicals) at 1:1000 overnight at 4 °C. The following day the cells were washed 3 times for 5 minutes each and then incubated with anti-mouse antibody labeled with AF647 (A21237, Invitrogen) for 1 hour at room temperature. Then the rabbit anti-lamin b1 antibody (ab16048, Abcam) was incubated with the samples overnight at 4 °C. The cells were washed with wash buffer (as above) 3 times for 5 minutes each and then anti-rabbit antibody labeled with CF568 (20099, Biotium) incubated with the samples for 1 hour at room temperature. Finally, the cells were washed with wash buffer for 3 times, 5 minutes each, and then rinsed.

#### 7.6 Osmostic Shock and Lamin and Coilin Immunofluorescence Sample Preparation

In large format experiments, U-2 OS cells were plated at a density of 5,000 to 10,000 cells per well for a final density of 20,000 cells per well the next day in clear Ibidi 96-well plate (#89626-C). For small format experiments, either HeLa or U2OS cells were plated at a density of 20,000 to 30,000 cells per well for a final density of 60,000 cells per well in an glass-bottomed Ibidi 8-well slide (#80807).

Osmotic shock treatments were applied by supplementing 2.5 M KCl (Sigma) to the corresponding cell media to a final concentration of up to 200 mM and replacing the media in the treated well for one hour. Wash out was carried out by replacing with normal media twice and allowing cells to recover for up to 180 minutes. Cells were then fixed with 4% PFA in PBS for 10 min. For conditions with no recovery from osmotic shock, 4% PFA was supplemented with 2.5 M KCl to the corresponding concentration. Samples were blocked and permeabilized with 3% BSA and 0.1% Triton X-100 for 15 min.

Primary antibody stain was performed overnight at 4 °C with anti-Lamin B1 (rabbit, Abcam #16048) and anti-coilin (mouse, Abcam #87913) each diluted 1:1000 in 3% BSA and 0.1% Triton X-100. Samples were washed three times with PBS and secondary stained for one hour at room temperature with anti-rabbit CF660C (goat, Biotium #20369) and anti-mouse Cy3B [Goat-anti-mouse antibodies (Jackson ImmunoResearch, 115-005-146 Lot:139375, 1.8 mg/mL) conjugated to Cy3B NHS ester (GE Healthcare Bio-Sciences Corp., PA63101) following the protocol used in [23]] each diluted 1:1000 in 3% BSA and 0.1% Triton X-100. Samples were washed three times with PBS and post fixed with 4% PFA for ten minutes.
